# Pathological microcircuits and epileptiform events in patient hippocampal slices

**DOI:** 10.1101/2024.11.13.623525

**Authors:** Matthew A.T. Elliott, John P. Andrews, Tjitse van der Molen, Jinghui Geng, Alex Spaeth, Anna Toledo, Kateryna Voitiuk, Cordero Core, Thomas Gillespie, Ari Sinervo, David F. Parks, Ash Robbins, Daniel Solís, Edward F. Chang, Tomasz Jan Nowakowski, Mircea Teodorescu, David Haussler, Tal Sharf

**Author notes:** Correspondence (T.S.). These authors contributed equally (Matthew A.T. Elliott, John P. Andrews).

## Abstract

How seizures begin at the level of microscopic neuronal circuits remains unknown. Advancements in high-density CMOS-based microelectrode arrays can be harnessed to study neuronal network activity with unprecedented spatial and temporal resolution. We use high-density electrophysiology recordings to probe the network activity of human hippocampal brain slices from six patients with mesial temporal lobe epilepsy. Two slices from the dentate gyrus exhibited epileptiform activity in the presence of low magnesium media with kainic acid. Both slices exhibit network oscillations indicative of a reciprocally connected circuit, which is unexpected under normal physiological conditions. Future studies may apply this approach to elucidate the network signals that underlie seizure initiation.

## Introduction

Mesial temporal lobe epilepsy is the most common form of drug refractory epilepsy in adults^1^. The cause of mesial temporal lobe epilepsy remains unknown despite decades of research on its basic mechanisms, spanning numerous animal models and the direct study of human brain slice physiology^2^. Studies often rely on histochemical surveys^3^, which lack functional context in describing neuronal behavior.

A limitation in prior work is the level of spatial and temporal resolution of the microcircuits implicated in epilepsy^4^. The recent advent of high-density CMOS-based microelectrode arrays allow for densely configured electrodes tiled across thousands of sites to record hundreds of networked neurons within mammalian brain slices^5–7^. This technology is now enabling clinicians to map neural dynamics of *in vivo* and *ex vivo* brain tissue at spatiotemporal scales previously inaccessible^8–11^.

In this study, we analyze circuit dynamics from the first experiments to collect large-scale electrophysiology recordings from patients suffering from mesial temporal lobe epilepsy using high-density microelectrode arrays^8^. Resected hippocampal tissue was sliced to 300*μ*m, incubated, and then plated onto a microelectrode array (**Fig. 1b**). In this work we analyzed six slices (S1-S6): four from the dentate gyrus, and two from CA1 (**Extended Data Table 1, Extended Data Fig. 1**)^12^. Two dentate gyrus slices from different patients exhibited seizure-like behavior after the administration of kainic acid. We observe a strikingly similar pattern of neuronal firing and local field potential (LFP) oscillations in the theta-band (4-8 Hz) that propagate as coherent waves during the onset of epileptiform bursting activity.

**Fig. 1:**
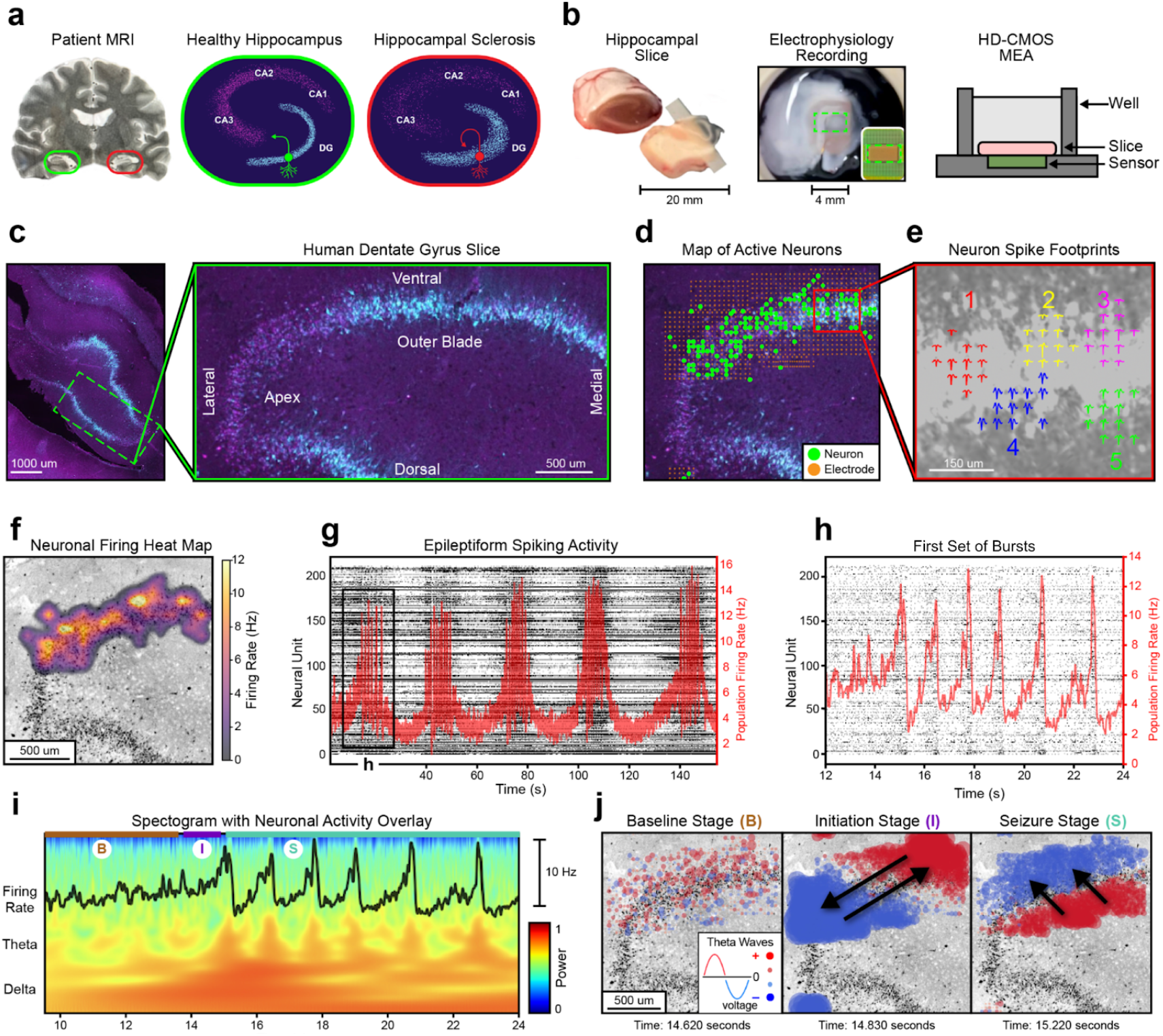
Spatially mapping epileptiform dynamics within the human dentate gyrus. **a,** Left: Patient MRI highlighting hippocampal sclerosis (red) compared to healthy tissue (green). Middle: Diagram of a coronal section from a healthy hippocampal slice. Primary regions are labeled (DG, CA1-3), and a depiction of a standard granule cell is displayed (green). Right: Diagram of sclerotic tissue. Exhibited pathologies include cellular loss, granule cell dispersion, and mossy fiber sprouting (red neuron). **b,** Resected patient hippocampal tissue is sliced to 300*μ*m and placed on an high-density microelectrode array (HD-CMOS MEA). A green rectangle depicts the region where electrophysiology data is recorded. **c,** NeuN (magenta) and eYFP (cyan) immunohistochemistry of a hippocampal slice transduced with AAV9-CAMK2A-HcKCR1-eYFP, with cyan representing CAMK2A expressing cell clustered in the granule cell layer of the dentate gyrus. Right: Magnified image of dentate gyrus, with subregions labeled. **d,** Spatial map of recording electrodes and neural units from the slice in **c,** discerned through spike sorting. **e,** Spatial footprints of five neurons action potentials from inset in **d.** For each neuron, an averaged 3ms action potential is displayed on top of its recording site. **f,** Heatmap visualization of neuronal firing rates collected across the duration of the recording from **d. g,** A raster plot of spike events shown as black dots (left axis) with the population average firing rate shown in red (right axis). Neural activity is from the first 150 seconds of epileptiform activity for the slice in **d. h,** A zoomed in raster of the initiation of bursting activity in **g. i,** A spectrogram at the initiation of bursting activity, highlighting delta and theta-band activity with the firing rate overlayed on top. Based on theta wave behavior, the recording is divided into three sections, baseline (B), initiation (I), and seizure (S) stage. **j,** Spatial plots of theta waves during baseline, initiation, and seizure stages. Red signifies a positive voltage, and blue signifies a negative voltage. Black arrows display the direction of wavefront propagations across time. See **Video 1** (https://youtu.be/sm3gKk-1Og4).

These results coincide with numerous proposed models of epileptogenesis^13,14^. Implicated biological mechanisms differ between model, and their role in pathogenesis is a topic of debate^15^. However, many consider hyperexcitation between granule cells in the dentate gyrus as the locus of seizure events. One well-recognized model is the mossy fiber sprouting hypothesis (**Fig. 1a**), which focuses on axonal outgrowths from granule cells forming new local connections in the dentate gyrus^16,17^. It postulates that this increased connectivity may establish a feedback loop of self-perpetuating hyperexcitation^18^. However, experimental evidence of hyperexcitability remains incomplete, due to an inability to map neuronal dysfunction at the microcircuit level.

We used high-resolution electrophysiology recordings to build anatomical maps of single unit neuronal activity overlaid onto histochemical stains from subregions of the hippocampus. Immunohistochemistry revealed CAMK2A-positve neurons clustered in the granule cell layer of the dentate gyrus (**Fig. 1c**). This is consistent with single nucleus sequencing data documenting high expression of CAMK2A in granule cells compared to other hippocampal neurons^19^. While slices expressed a channelrodopsin HcKCR1-eYFP fusion protein driven by a CAMK2A promoter (AAV9-CAMK2A-HcKCR1-eYFP)^20^, optogenetic results are discussed in Andrews *et al*.^8^.

To visualize the electrophysiologic data, we project the locations of neurons and electrodes onto a 2D map. The locations of each recording electrode and neuronal unit, discerned through spike sorting, were overlaid onto the histology images by best approximation for all six slices (**Fig. 1d, Extended Data Fig. 2**)^21^. The high-density electrophysiology recordings provide resolution at the level of an individual neuron’s extracellular footprint, allowing for spatial plots and subsequent analysis of single neuron behavior across large distributed networks within the human hippocampus (**Fig. 1e-f**)^9,22,23^.

## Results

We began our analysis by assessing whether epileptiform behavior was present in our hippocampal slices. After administering kainic acid to slice S1, synchronized rhythmic bursting activity was observed (**Fig. 1G**). Kainic acid is a widely used convulsant that induces seizure-like activity by activating kainate receptors, leading to sustained neuronal hyperexcitability^24^. Visualization of the onset of burst synchronization reveals a dynamical substructure that contains approximately 6-8 bursts grouped within larger burst epochs (**Fig. 1h**)^25^. A spectrogram from S1 displays the relative power of the local field potential (LFP) from a representative electrode at the initiation of bursting (**Fig. 1i**). We observe electrophysiological characteristics consistent with seizure-like events after administering kainic acid. These include a defined increase in the amplitude of theta-band oscillations, coherence between theta-band activity and bursting behavior, and a large upwelling in the delta-band frequency at epileptiform onset^26–28^. These characteristics remained present across different electrode sites where the LFP traveled as a coherent wave (**Extended Data Fig. 3**). Epileptiform behavior was also present when kainic acid was administered to tissue from a different patient with mesial temporal lobe epilepsy (**Extended Data Fig. 5**). To understand what might initiate seizure-like events, we investigated theta-band oscillations at the onset of rhythmic bursting activity. We projected the theta-band voltage signals from approximately 1000 electrodes onto the slice histology (**Fig. 1j**) in order to visualize the high-density electrophysiologic data while preserving its spatial dimension as well as its time dimension. In these plots red represents positive voltage and blue represents negative voltage. The changing dynamics of the theta wave propagations are best observed by watching **Video 1** (https://youtu.be/sm3gKk-1Og4).

We observed theta-band propagations at the onset of seizure-like behavior that transitions between two orthogonal modes. The onset of bursting activity following administration of kainic acid (**Fig. 1i**) was divided into three stages that we distinguish based on bursting activity. We refer to these stages as “baseline” (prior to burst activity), “initiation” (the first burst), and “seizure” (subsequent bursts)^29,30^ (**Fig. 1j, Video 1**). At the stage we term baseline, there is no discernable pattern of theta propagations. The initiation stage occurs at the transition to seizure-like activity, where the amplitude of theta wave oscillations exceed background noise^31^. During the initiation stage, theta propagations form a standing wave that oscillates across the length of the granule cell layer (**Extended Data Fig. 4d**). In this stage, theta oscillations emerge and travel across the axis of the granule cell layer (**Fig. 1j**). The direction these waves propagate is unique to the beginning of bursting activity. (**Video 1-4**). Following initiation and during all subsequent bursts, the theta oscillations form a rolling wave orthogonal to the direction observed during the initiation stage, moving along the granule cell layer from the hilar to outer aspect (**Extended Data Fig. 4e**). We refer to this stereotyped wave activity during subsequent bursts as the seizure stage. The same stages of baseline, initiation, and seizure were observed when kainic acid was administered to a second slice from the dentate gyrus of a different patient (S2) (**Extended Data Fig. 5**).

We checked that observed coherence in theta-band propagations was unique to epileptiform activity. For each stage of the recording, electrodes were clustered based on their average time delay between theta propagations (**Extended Data Fig. 4g**). With this technique, clusterings present the directionality of the dominant wavefront for each stage. Spatial maps of the electrode clusterings replicated the behavior seen in theta wave propagations (**Extended Data Fig. 4h**). The slices with no epileptiform activity (S3-S6) did not contain spatially coherent theta-band wave propagation patterns (**Extended Data Fig. 6**).

We next considered if the theta-band propagations observed at the initiation stage might reflect a distinct spatiotemporal pattern of neuronal activity. We constructed spatial plots of changes in the single unit firing rate and compared them to the timing of theta propagations. We found that theta-band oscillations during the initiation stage of S1 are temporally aligned with spatiotemporal oscillations in single unit spiking activity, suggesting some form of recurrent feedback at epileptiform onset^18^. Neuronal activity from the first burst of S1 was divided into three sub-bursts (**Fig. 2a**). The peaks and troughs of sub-bursts aligned with when the two modes in theta-band activity were maximally distinct from each other. Calculating the change in single unit firing from sub-burst to sub-burst produced heatmaps with similar spatiotemporal oscillations to those seen in theta-band propagations. We observed a similar pattern of activity in a separate slice (S2) of the dentate gyrus from a different patient, suggesting a conserved feedback loop drives epileptiform initiation (**Extended Data Fig. 5f**). A different pattern of coherence between theta waves and spiking behavior occurred during the seizure stage (**Extended Data Fig. 7, Video 3-4**). Taken together, these findings support previous *in vivo* work linking local neuronal activity to LFP oscillations^32,33^ and provide an approach to link macroscale LFP dynamics to neuronal dynamics at the microcircuit level.

**Fig. 2:**
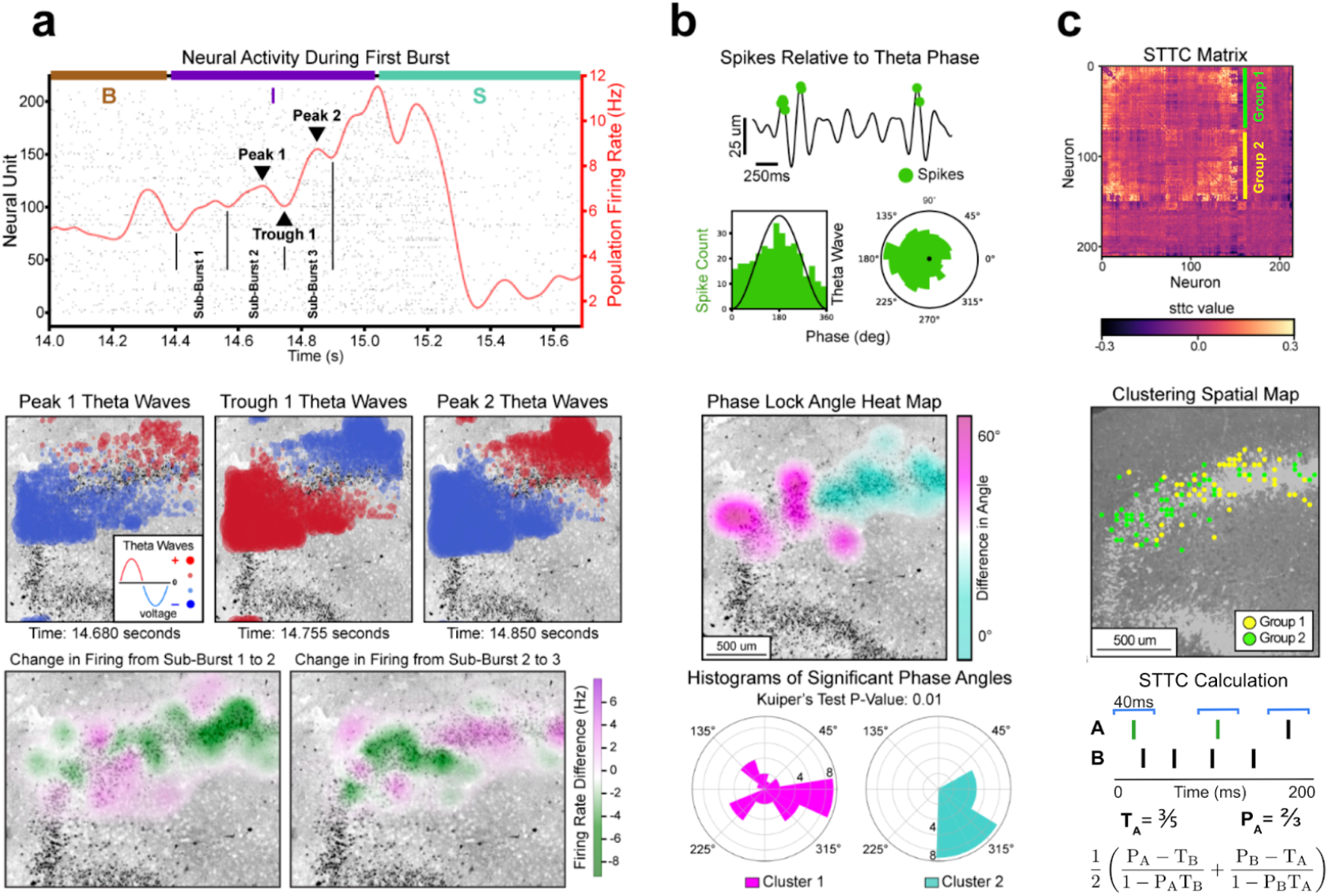
Recurrent network oscillations at onset of seizure-like behavior. **a,** Top: Neural activity from the first burst of the recording with baseline, initiation, and seizure phases labeled by B, I, and S, respectively. In the initiation phase, sub-bursts, peaks, and troughs are labeled. Middle: Spatial plot of theta wave activity taken at the time points of the labeled peaks and troughs. Bottom: Heatmap of the change in firing from sub-burst to sub-burst. During initiation, firing activity oscillates between the two modes observed during theta wave activity. **b,** Top: Example of a neural unit that is significantly phase locked (Rayleigh p-value<0.05). Spike events are projected to their corresponding location on a theta wave. Two histograms, polar and nonpolar, display the frequency of spikes based on theta phase angle. Middle: Heatmap of the average difference in phase angles between all significantly phase locked units across the entire recording. Bottom: Polar histograms of neural phase angles from the two clusters in the heatmap above. Clusters have significantly different angular distributions (Kuiper’s p-value<0.01) **c,** Top: The spike time tiling coefficient (STTC) matrix, with units organized based on agglomerative hierarchical clustering. Green and yellow squares indicate groupings observed from clustering. Middle: Spatial plot of neurons based on their grouping from the STTC matrix. Bottom: Schematic of STTC calculation for neurons A and B, presenting raster and tiling windows for A. The equation is shown at the bottom. T_A_ is the proportion of the recording spanned by A’s tiling windows. P_A_ is the proportion of A’s spikes that contain a spike from B in their tiling window.

After analyzing neuronal dynamics during epileptiform onset, we next considered if these dynamics may be part of a larger pattern of neuronal behavior present across the entire length of the recording. Such a pattern might suggest an underlying mechanism related to the dynamics observed in the initiation stage. Phase locking is a common tool for measuring the synchronization between neural spiking behavior and theta rhythms^34,35^ in the human hippocampus^36^. Phase locking analysis across the entire recording provided results consistent with those seen in the initiation stage (**Fig. 2b**). Significantly phase locked units were determined by considering the correspondence between their spike times and theta-band phase (Rayleigh p-value < 0.05)^37^. A heatmap of the difference in phase angles between significant units produced a bimodal plot similar to those previously shown (**Fig. 2b middle**). Clustering based on the two modes of significant units yielded two groups with significantly different phase angles (Kuiper’s p-value = 0.01). When evaluating the phase locking dynamics across all slices, we observed considerably more phase locked units in the two slices with epileptiform activity, with phase locking particularly high during seizure-like events (**Extended Data Fig. 8, Extended Data Table 2**).

Spiking behavior across the entire recording recapitulates the spatial dynamics observed in phase locking (**Fig. 2c**). Two groupings of neuronal populations were created by performing hierarchical clustering on the spike time tiling coefficient (STTC) matrix, a more robust measure of correlations between neuronal spike trains that are often confounded by firing rate^38^. A spatial map of the neuronal clusters divides the granule cell layer into the same two modes seen in previous plots. Similar results were observed in the second epileptiform slice (S2) (**Extended Data Fig. 5h**). A second clustering method based on the eigendecomposition of the STTC matrix reproduced these results (**Extended Data Fig. 9**).

We noticed a marked resemblance in the spatial plots from the initiation stage (**Fig 2a**) when compared to the spatial plots from our whole recording analysis (**Fig 2b-c**), with all plots dividing the granule cell layer into two modes along its long axis. This led us to ask if our high neuronal resolution dataset could be used to reveal anatomical organization that might explain recurrent behavior. We constructed an *in silico* model to simulate functional connectivity within the dentate gyrus (**Extended Data Fig. 12**)^39^ and found that increasing pairwise interactions between granule cells produced seizure-like bursting events. This led us to consider if constructing anatomical diagrams between neuronal pairs might elucidate anatomically distinct functional correlations within our slices.

We created neuronal circuit diagrams to display the direction of neuronal spiking signal propagations across the tissue as a vector arrow plot. The circuit diagram for S1 is created using the entire recording across baseline, initiation, and seizure stages (**Fig. 3a**). The arrows in the circuit diagram are derived from correlations between neuronal pairs measured across 30 ms latency windows^40^ (**Fig. 3a left**), with the color denoted by the angle between each pair. This time window was used to infer functional correlations on timescales that reflect polysynaptic transmission across the slice^41^, rather than a direct measure of monosynaptic spike transmission probabilities, which were not observed in our data^42,43^. The circuit diagram for slice S1 displays two clusters of arrows (green and red), consistent with the bimodal activity we observed in phase locking and neuronal firing analysis (**Fig. 2a,b**). The angular distribution of the circuit diagram reveals two reciprocally connected populations segregated in space. To quantify this, we created a pairwise correlation angle histogram (bottom-right of **Fig. 3a**) that plots the total number of correlated spike events by propagation angle. For slice S1, the polar histogram shows bipolar angular pairwise correlations that are 180° out of phase and aligned with the propagation of theta waves during epileptiform initiation events.

**Fig. 3:**
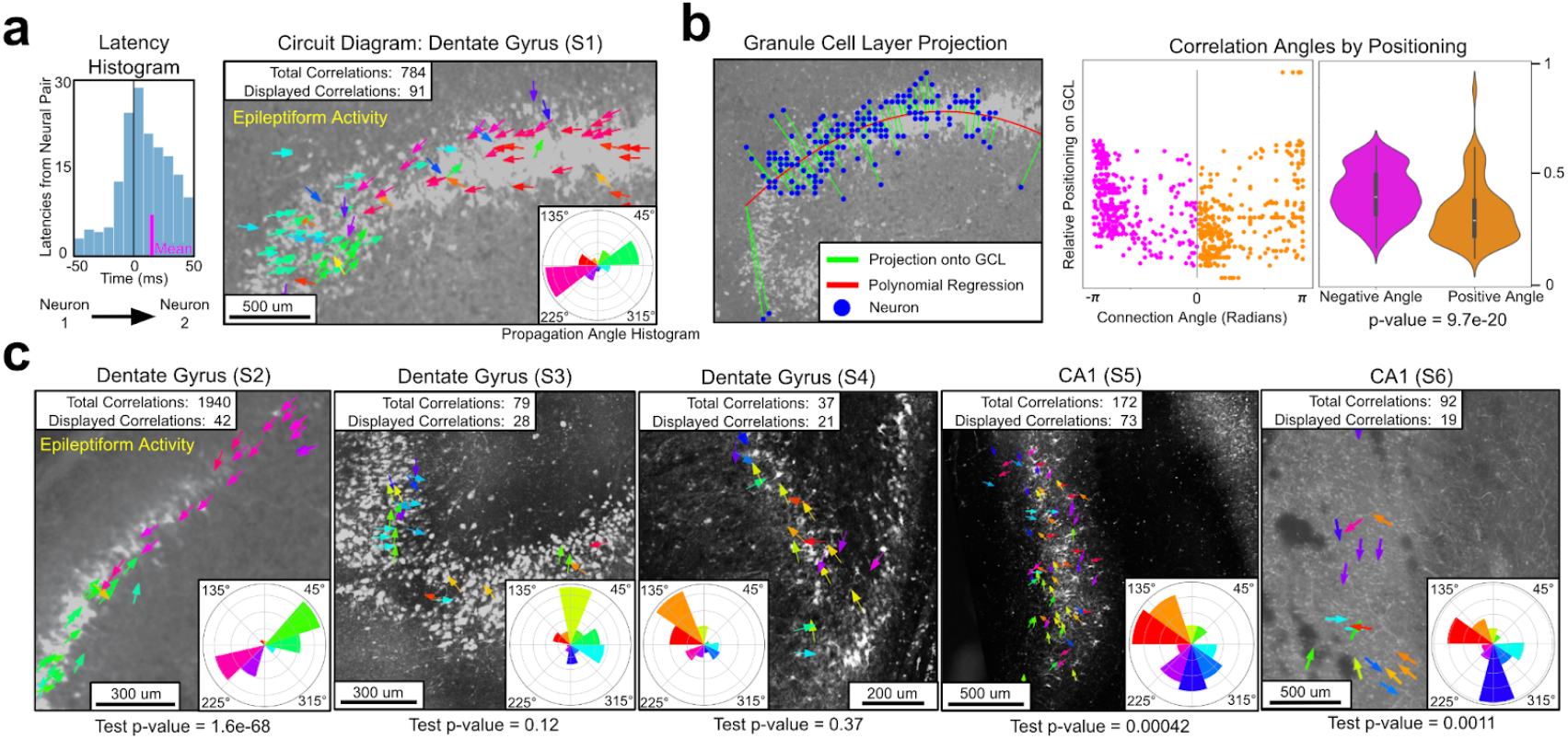
Reciprocal pairwise neuronal correlations are unique to epileptiform slices. **a,** Left: A histogram of spike time latencies between two neurons with a non-zero mean used to construct a directional signal propagation vector (t-test p-value<1e-10, mean=10ms). Right: A neuronal circuit diagram displaying 91 of the 784 significant functional correlations. Correlations are colored based on the angle spikes propagate between neurons. The propagation angle histogram (bottom-right inset) displays the angular frequencies from all spike propagation events. The pairwise correlation angles are bipolar, highlighting the slice’s recurrent functional dynamics. **b,** Spatial test of reciprocal correlation significance along the curvilinear axis of the granular cell layer (GCL). Left: The relative placement (green line) of every neuron in S1 with respect to the GCL, as approximated by a second order polynomial regression (red line) fitted to the neurons (blue dots). Right: A scatter plot from S1 compares every propagation arrow’s angle (in radians) to its relative positioning on the GCL. Data is grouped based on whether the angle is above or below zero (orange and magenta). A two-tailed t-test determined the two groups are significantly different (p-value<1e-19) **c,**Directional diagrams for every slice (S2-S6). The second slice with epileptiform activity (S2) exhibits the same bipolar angular correlations along the GCL observed in **a.** The epileptiform slices also have considerably lower p-values from the spatial test in **b.** Using the propagation angle histograms, we observe that the other slices from the dentate gyrus (S3-S4) have unidirectional circuits, and slices from CA1 have bimodal, but not reciprocally distributed correlation angles.

We compared the circuit diagrams and the propagation angle histograms from all eight recordings and found a microcircuit, derived for pairwise spike train correlations, that was unique to the epileptiform slices (**Fig. 3c**). The second slice with epileptiform activity (S2) also had a bimodal circuit diagram, with spikes propagating toward each other, and a bipolar angle histogram (see inset of **Fig. 3a**). This is consistent with the recurrent behavior observed at seizure-like initiation for S2. For both S1 and S2, the non-epileptiform recordings prior to administering kainic acid had the same circuit diagram as during epileptiform activity (**Extended Data Fig. 10**). This suggests that the bimodal circuit of S1 and S2 is a structural quality of the slices and not a by-product, only present during seizure-like bursting activity.

None of the non-epileptiform slices displayed the bimodal circuit diagram observed from the epileptiform slices. Both of the non-epileptiform dentate gyrus slices (S3 and S4) had a unimodal circuit diagram, with spike signals propagating away from the apex. Notably, slice S4 was administered kainic acid, but did not exhibit any epileptiform behavior or the circuit dynamics of S1 and S2. The slices from CA1 (S5 and S6) had evenly distributed circuit diagrams and a bimodal, but not bipolar, with spike signals oriented along the perforant pathway. In the case of all recordings, slices from the same subregion followed similar circuit behavior. A statistical test was performed to check if the differences we observed in circuit diagrams were due to bias in the geometric layout of neurons (**Fig. 3b**). The test accounts for the angles that spike signals propagate relative to their positioning along the granule cell layer. If the bipolar plots from epileptiform slices were produced by bias in neuronal geometry, the test’s p-values would be similar for all slices, however, we found the epileptiform slices to be orders of magnitude more significant (**Extended Data Table 3**).

## Discussion

In summary, using high resolution CMOS-based microelectrode array recordings, we relate the spatial dynamics of signal oscillations at the onset of seizure-like activity (**Fig. 2**) to circuit level network behavior in human hippocampal slices (**Fig. 3**). Two hippocampal slices from different epilepsy patients display bimodal neuronal activity patterns that suggest recurrent circuits in the dentate gyrus may drive epileptiform activity (**Extended Data Fig. 4**,**5**,**8**). Furthermore, recurrent oscillations in both theta-band LFPs and neural firing activity were observed between two spatially distinct activity modes at the initiation of epileptiform behavior. These findings are consistent with simulations demonstrating that increased granule cell interconnectivity in the dentate gyrus leads to hyperexcitation and seizure-like events (**Extended Data Fig. 12**).

This study is not without limitations. The availability of surgical samples is a restricting factor for these experiments. All slices are derived from patients with refractory epilepsy, therefore there is no true control group for comparison. However, we observed that not all slices produced seizure-like activity when provoked with low-magnesium and kainic acid media. Thus, in lieu of true controls, we compare slices that elicited seizure-like activity upon chemical provocation to those that did not. Moreover, this is a retrospective analysis of the first experiments to use high-density CMOS-based microelectrode arrays on human brain slices, with experimental design geared toward testing optogenetic interventions^8^. Confounding factors are introduced by the original experiment. Notably, organotypic slices were used, instead of acute, and optogenetic inhibition was performed on the latter portion of the kainic acid recordings (after epileptiform initiation). While it is possible that expression of the HcKCR1-eYFP protein could affect neuronal physiology even in the absence of activation by light illumination, the same critique could be levied at most essential optogenetic physiology literature.

In conclusion, recent advancements in high-density microelectrode arrays allow for the study of large populations of neurons with single cell resolution, presenting the opportunity to examine models of mesoscale network behavior. When applied to mesial temporal lobe epilepsy, our data supports recurrent communication between granule cells in the dentate gyrus as a possible mechanism of excitation and seizure initiation. These results are consistent with numerous biological models of epileptogenesis, including the mossy fiber sprouting hypothesis, which considers hyperexcitation in the dentate gyrus as the locus of disease^16,17^. By integrating these computational methods with novel biological approaches for elucidating microscopic neural circuits^44^, larger prospective studies have the potential to localize neurological mechanisms underpinning epilepsy. A more complete understanding of seizure circuitry may pave the way for new therapeutic approaches^45,46^.

## Methods

### Tissue preparation

Samples were collected from patients undergoing temporal lobectomy with hippocampectomy for refractory epilepsy. We obtained signed patient consent and approval from the University of California-San Francisco Institutional Review Board. Informed consent from patients was obtained prior to surgical resections. Patient characteristics are detailed in Extended Data Table 4.

Tissue transport, preparation, and culture were adapted from prior studies^8,47^. To briefly summarize from Andrew et al., tissue was collected from the operating room, sliced into 300µM sections using a vibratome, and put in sterile artificial cerebrospinal fluid (aCSF) bubbled with carbogen. The pH of the aCSF was titrated to 7.3–7.4 with hydrochloric acid and had an osmolality of 300–305 mOsmoles/Kg. Slices were plated on cell-culture inserts at the air-liquid interface and transduced with a CAMK2A promoter via an adeno-associated virus. Slices briefly recovered in aCSF warmed to 33°C before plating on cell culture inserts at the air-liquid interface and incubated at 37°C in 5% CO2 incubators. Slices were incubated for 4-8 days.

On the day of the recording, slices were incubated for one hour with Matrigel and then plated on high-density microelectrode arrays with minimal culture media during the recording. Slices were floated into the MEA wells and then media was aspirated slowly such that the slice descended onto the recording surface. A harp was placed on the slice to press it against the recording array. For slices with low magnesium media, the physiologic media was made without any magnesium (MgSO4) added. For slices with Kainic acid experiments (S1, S2, and S4), 100nM kainic acid (KA) was dripped directly onto the slice. A continuous recording was taken during the administration of KA, with epileptiform behavior being observed in S1 and S2 within 30 seconds. Slices that did not show electrophysiologic evidence of at least 50 active neural units were excluded from electrophysiologic analyses.

### Experimental design, reproducibility, and inclusion/exclusion criteria

The sample size was maximized based on the availability of human brain tissue. No statistical method was used to predetermine sample size, but our sizes are similar to those in previous publications^8,48,49^. Experiments were run in the order that tissue became available. No randomization was performed. Data collection and analysis were not performed blind to the conditions of the experiments. All the slices that were analyzed were from adult patients with refractory epilepsy. The regions of the hippocampus from which recordings were taken were selected based on where tissue appeared healthiest.

We required higher levels of neural activity than what was necessary for the original study, thus we excluded any slice with less than 50 neurons after spike sorting. Of the 12 slices from the original study, six were analyzed. For transparency, results from all 6 slices are present in our analysis. See Extended Data Fig. 1,2,10 and Extended Data Tables 1-3. The main figures display results for the primary slice with epileptiform activity, S1. Corresponding results from the second slice with epileptiform activity, S2, are in Extended Data Fig. 5. Results for the four non-epileptiform slices (S3-6) are in Extended Data Fig. 6.

### Immunohistochemistry

For the histology images shown in Fig. 1c-d and Extended Data Fig. 1, the following antibodies were used for immunohistochemistry. Slices were infected with the AAV9-CAMK2A-HcKCR1-eYFP adeno-associated viral vector prior to staining, which utilized anti-NeuN (magenta), demonstrating dense staining of CAMK2A-expressing neurons clustered in the granule cell layer of the dentate gyrus (**Fig. 1c**).

NeuN: guinea pig anti-NeuN, Millipore, ABN90, dilution 1:1000, lot#4077530

### Data acquisition and spike sorting

Extracellular field potentials were sampled at 20kHz from up to 1,024 electrodes using an high-density CMOS microelectrode array technology (MaxOne, Maxwell Biosystems, Zurich, Switzerland)^50^. The array contains 26,400 recording electrodes with a diameter of 7.5μm at a center-to-center distance of 17.5 μm. At the beginning of the experiment, an activity scan assay was performed across all electrodes. Approximately 1,000 recording electrodes were manually selected based on the most active regions found in the scan. After the experiment, raw activity data was saved to an HDF5 file on local memory.

Raw extracellular recordings were bandpass filtered between 300-6000Hz and then spike sorted in Kilosort2^51^ to extract single neural unit locations and activity. Sorting was performed on the Pacific Research Platform computing cluster^52^. Kilosort2’s results were manually curated using Phy GUI^53^ by experienced researchers who took into consideration each unit’s spike waveform, correlogram, and interspike interval violations.

### Spatially mapping electrodes and neurons

Fig. 1d and Extended Data Fig. 2 display the spatial locations of recording electrodes and neural units mapped to histology images. After a slice was plated onto the microelectrode array, an upright microscope (MS08B, Dino-Lite) photographed the tissue. We mapped neurons and electrodes by comparing the microscopy image to the histology. Specific locations of electrodes were extracted from the H5 file produced by the recording. The placement of neural units was provided by the Kilosort 2 spike sorting algorithm.

### Spike rasters with population level firing activity

Fig. 1g-h and Fig. 2a contain neural spike rasters with the population level firing rate overlayed on top. The population firing rate is calculated by, first, summing the total number of spikes in each millisecond bin. A moving average of the spike counts is created by applying a 1D Gaussian filter to the bins, with a standard deviation between 10-20ms. Each 1ms bin of spike counts is divided by the total number of neurons and multiplied by 1000 in order to correspond to the standard formula for firing rate.

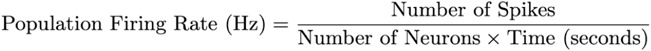

### Neuronal Firing Activity Heatmaps

A spatial heatmap of firing activity is displayed in Fig. 1f, illustrating the average neuronal firing rate (in Hz) across the granule cell layer. First, the average firing rate for each neuron was calculated by dividing the total number of spikes by the recording duration. Neurons were spatially mapped into a grid of 900 squares, each measuring approximately 58.3 µm x 58.3 µm. The average firing rate for each square was calculated based on the neurons it contained. A 2D Gaussian filter was applied to the grid in order to smooth the spatial distribution of firing rates. The filter replaces each point with a weighted average of its neighboring points. These weights are determined by a 2D Gaussian distribution, with a standard deviation of 37.9 µm. Fig. 2f and Extended Data Fig. 5f displays heatmaps of the difference in firing activity between neuronal sub-bursts using a method analogous to the one described above.

### Neuron Spatial footprints

Fig. 1e displays individual neurons’ spatial footprints. Kilosort 2 saves the putative footprints for each neural unit found through spike sorting. Representative electrodes are detected for each neural unit by averaging the amplitudes across spikes and choosing the 12 channels with maximum amplitude. The waveforms displayed are an average of the action potentials that occurred for that electrode.

### Spectrogram

Fig. 1i and Extended Data Fig. 3 display spectrograms. The spectral analysis shows the signal strength of different subbands in the local field potential of a single electrode. We first filter the raw voltage signal with a bandpass filter between 0.1-100 Hz. Signals are downsampled from 20 kHz to 1 kHz. A second bandpass filter is then applied to extract the sub-band frequencies of delta (0.5 -4 Hz), theta (4 -8 Hz), alpha (8 -13 Hz), beta (13 -30 Hz) and gamma (30 -50 Hz) waves. To plot the spectrogram, we run a continuous wavelet transform on the local field potential data. This transform is performed using the complex Morlet wavelet (‘cmor1-1’ in pycwt), which computes wavelet coefficients and corresponding frequencies. The power is computed as the magnitude squared of the wavelet coefficients and then smoothed using a Gaussian filter (sigma=2).

### Theta wave activity plots and videos

Theta wave activity plots provide a spatial map of the theta wave values from all, roughly, 1000 electrodes at a single cross-section in time (Fig. 1j and Extended Data Fig. 4-5). A standard neuroscience protocol was used to calculate the theta wave values for each electrode. A fourth order Butterworth bandpass filter selecting frequencies between 4-8Hz was applied to the raw voltage data from each electrode. For visualization in the spatial plots, resulting theta waves are normalized by electrode, with values between [-1, 1]. The theta value from each electrode is represented by a circle, centered at the spatial location of the electrode. The color of the circle is red if the voltage of the electrode’s theta wave value is positive, and blue if it is negative. The size of the circle scales with the absolute amplitude of the voltage, making circles with a higher magnitude larger.

Every theta wave activity plot has a corresponding video, which provides a clearer understanding of how theta wave propagations evolve through time (Extended Data Videos 1-4). Videos of theta wave activity move at a pace 10-20 times slower than real time. The frames in the video change at a 5ms time interval. The video displays theta activity on the left and neural firing activity on the right. Having these plots side-by-side is useful for understanding the coherence between theta activity and population level firing activity (see Video 3, Extended Data Fig. 7).

### Electrode spatial clustering algorithm

For slices with epileptiform activity (S1 and S2), theta propagations observed in the baseline, initiation, and seizure phases (Video 1) were verified using a spatial clustering algorithm (Extended Data Fig. 4g and 5e). The clusterings resemble the theta activity seen for the corresponding phase. Electrodes were clustered based on the average time delay (lag time) between theta waves as they propagated across the electrodes.Cross-correlation analysis was performed between all pairs of electrodes (scipy function: correlate). The lag time (ms) that maximized the absolute correlation between a pair of electrodes was selected as the pair’s time delay. Only lag times between [-40,40] ms were considered. A square matrix is created using the lag times from all electrode pairs (Extended Data Fig. 4f). K-means clustering is performed on the lag times matrix using an N of two clusters. The spatial clusterings of the electrodes are then observed by plotting electrodes by location (Extended Data Fig 4g).

### Phase locking neurons to theta wave activity

Phase locking analysis was performed in Fig. 2b, Extended Data Fig. 8, and Extended Data Table 2^54^. First, the phase angles of the LFP filtered in the theta frequency band (4-8Hz) were obtained by a standard Hilbert transformation on the time series and subsequently calculating the angle between the real and imaginary components. For each spike sorted unit, the upper envelope of the theta filtered LFP was also calculated. For all spikes that occurred while this upper envelope was above the RMS of the whole time series, the theta phase angle of the spikes were stored. The Rayleigh criterion was used to test the non-uniformity of the phase angles over all the selected spikes (0°, 360°). Spikes were considered to be phase-locked to theta if they passed the Rayleigh criteria test for non-uniformity (p < 0.05). This was done separately for seizure and non-seizure phases in the recording (Extended Data Fig. 8). For each significantly phase-locked unit, the circular mean was computed over all selected spikes to obtain the average phase locked angle. The Rayleigh criteria test for non-uniformity is the polar analog to the one-sample T-test. Its R-statistic (analog to t-statistic) is calculated as follows:

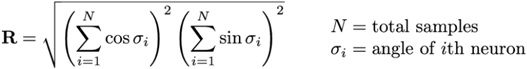

### Methodology for heatmap of phase locked angles

Heatmaps presenting the spatial difference in phase locked angles across the tissue were presented for slices S1 and S2 (Fig. 2b and Extended Data Fig. 5g). Heatmaps were constructed using only neurons that were significantly phase locked (Rayleigh p-value<0.05). The polar mean was used to find the primary phase angle each neuron was phase locked to. A polar histogram of all significant phase angles was constructed to determine the most frequently occurring angular direction (polar mode). The absolute angular difference from the polar mode (L1 norm) was calculated. These differences were used to construct a spatial heatmap of polar angles.

Given the angular values of each neuron, the methodology for constructing the heatmap is similar to the heatmaps for firing activity described above. Neurons’ angular values were mapped to a grid of squares, with the value of each square calculated based on the average of the neurons it contained. Then a 2D Gaussian filter was used to construct a smoothed spatial distribution, using a standard deviation of 70µm. The polar histograms next to the heatmap display angular frequencies corresponding to the two clusters observed from the heatmap. They are created by partitioning the phase locked neurons based on their angular difference and then plotting the resulting clusters. The Kuiper test was used to compare the angular distributions between the two clusters.

### Methodology for STTC hierarchical clustering of neurons

For both epileptiform recordings (S1 and S2), we provide spatial plots of neural clusters (Fig. 2c and Extended Data Fig. 5h). To create the clusters, a standard agglomerative hierarchical clustering algorithm was implemented on the spike time tiling coefficient (STTC) matrix. Because agglomerative clustering is sensitive to outliers, the STTC matrix’s maximum value was thresholded to 0.3 (roughly the 98th percentile). Clustering was done using the scipy.cluster.hierarchy package. The Euclidean pairwise distances between all pairs in the STTC matrix were calculated. Agglomerative hierarchical clustering was done on the pairwise distance to construct a linkage matrix corresponding to a dendrogram. The resulting hierarchical clustering tree was reordered to reflect the optimal leaf arrangement. The STTC matrix was reordered based on the optimal leaf ordering (Fig. 2c). Two clusters were observed in the resulting matrix. The neurons from these clusters were plotted spatially.

### Definition of circuit diagram propagation vector

In the circuit diagram figures (Fig. 2d and Extended Data Fig. 10), an individual arrow is defined as a circuit propagation vector. It represents a significant pairwise correlation between two neurons, where the base of the arrow is the location of the neuron from which the spike emanates and the angle of the arrow points toward the neuron that spikes afterward. Arrows are drawn at a constant fixed length, usually shorter than the distance between neural pairs. This is done to declutter the diagram.

Pairwise correlation vectors between neurons are selected by considering the latency events between every pair of neurons in the dataset^39^. Fig. 2d illustrates the methodology used to select pairwise correlations. Given a pair of neurons (n_1_,n_2_), for every spike that occurs in n_1_, we find the nearest occurring spike from n_2_, using the absolute difference in time as our metric. We construct a latency distribution by considering events that occur within a window of [-30ms,30ms]. A two-tailed t-test is performed on the latency distribution to determine if it is significantly different from zero (p-value < 0.05). An arrow is created for all significant pairs. Of the 45,000 neuronal pairs of neurons in slice S1, 784 pairs were classified as significant.

A standardized protocol constructed circuit diagrams for all recordings (Extended Data Fig. 10). To reduce computation time, only neural pairs with a spike time tiling coefficient value greater than 0.01 were considered. Pairs had their latency distributions calculated. Latency distributions with less than 25 latency events or with an absolute mean latency less than 1ms were disregarded. A two-tailed t-test was performed on all remaining pairs. If the pair’s p-value was significant (<0.05), the neuronal pair is included as a pairwise correlation in the circuit diagram.

### Displaying simplified circuit diagrams

The neural circuit diagrams in Fig. 3 and Extended Data Fig. 10 display only a fraction of all arrows representing significant pairwise correlations. Displaying all arrows would make the plot difficult to interpret. Extended Data Fig. 11 illustrates the protocol used to select a representative subsample of vectors.

First, we aggregate propagation vectors based on the neuron that is propagating the signal. Multiple connections emanating from a neuron are replaced with a single arrow. The polar mean (scipy: circmean) is used to aggregate the angles of all emanating pairwise correlations, with each vector weighted by its number of latency events. Aggregating propagation vectors sometimes display an arrow that is not representative of the original directions (see Extend Data Fig. 11). Vectors pointing in opposite directions are averaged to form a new arrow, not representative of either original pairwise correlation. Nonrepresentative aggregated arrows are removed by only considering aggregated connections whose standard deviation is below 0.5. The weighted polar standard deviation is used (scipy: circstd), with weights based on latency events. The remaining vectors are displayed in the circuit diagrams.

### Propagation angle histogram

At the bottom of every circuit diagram is an inset displaying the pairwise correlation angle histogram (Extended Data Fig. 10). The angle histogram displays the angular frequency of spike propagations across all circuit connections. The histogram is calculated over all significant pairwise correlations, not just the displayed vectors (see above). Pairwise propagations are weighted based on the number of latency events that occur within each pair. Colors in the angle histogram correspond with the color of arrows in the circuit diagram.

### Statistical test of circuit geometry

A statistical test was performed to check whether or not the recurrent circuits seen in the epileptiform slices (S1 and S2) are due to a bias caused by the geometry of neuronal locations. For all recordings, the spatial layout of neurons roughly followed a line across the granule cell layer (Slice 1-4) or pyramidal cell layer (Slice 5-6) (Extended Data Fig. 2). This test checks whether or not the relative location of neurons is the determining factor in the propagation angle or pairwise correlations. Fig. 3b provides an illustrative schematic of the test.

To find the relative positioning between neurons in a recording, they were first projected onto a line that goes through the center of the cell layer. The line was approximated by fitting a polynomial regression of degree two to all neurons. Neural positions were normalized with the leftmost position along the curve being 0.0 and the rightmost being 1.0. A scatter plot was created comparing the relative position of neuronal pairwise correlations to their corresponding angle (in Radians). All significant correlations were divided into two groups based on whether the connection angle was positive or negative. A two-tailed t-test was performed on the neural positionings of the two groups to test the extent to which they were significantly different.

If the observed circuit diagrams were due to bias in the neural geometry, the test’s p-values would be roughly similar for all slices, because relative neural positionings would be the determining factor in circuit angle. However, we found the bipolar recurrent circuit recordings (S1 and S2) to be orders of magnitude more significant, verifying that the phenomena we observe is not due to bias in neural geometry (Extended Data Table 3).

### Eigendecomposition for neural spatial clustering

We perform an eigendecomposition on the STTC matrix to spatially cluster neurons for slices with epileptiform activity (S1 and S2). An illustrative schematic of the methodology and its results are displayed in Extended Data Fig. 9. First, the STTC matrix was calculated. The STTC matrix is analogous to the commonly used correlation matrix, but has been shown to perform more robustly on neural data^38^. For our data, the STTC matrix had higher eigenvalues and a lower reconstruction error when compared to correlation. Like the correlation matrix, the STTC matrix is positive semidefinite, which means its eigendecomposition can be mathematically interpreted similarly to that of Principal Components Analysis (PCA). We plot the values from the first eigenvector spatially by coloring each neuron based on its eigenvector value. This results in a gradient that travels across the granule cell layer, similar to the bimodal clusters observed from hierarchical clustering (Figure 2c).

### Simulation of epileptiform behavior

We created a dynamical simulation of a hippocampal network as a proof of concept. Our model is a simplified version of a previously published *in silico* model of the human dentate gyrus which has been used to study disease progression in epilepsy^55^. Simulated connections between granule cells in the dentate gyrus were added as a variable fraction of the total cells within this model in order to probe its effect on seizure-like events. For further details, see the “Simulation” directory in the Github repository.

## Code Availability

All coding was done in Python, 3.10. The code used in the final processing and analysis of data is publicly available on GitHub. The repository has been integrated with GitHub Codespaces. Researchers can easily launch an online environment to reimplement our analysis. Instructions are in the GitHub repository: https://github.com/braingeneers/human_hippocampus

## Data Availability

All data used in the analysis and creation of figures is available in the GitHub repository: https://github.com/braingeneers/human_hippocampus

The repository includes neural spiking data, histology images, portions of raw electrophysiological data, and plots created for figures. The complete dataset of raw electrophysiological recordings from experiments is available on the DANDI public server: https://dandiarchive.org/dandiset/001132

## Acknowledgements

This work was supported by the Schmidt Futures Foundation SF 857 and the National Human Genome Research Institute under Award number 1RM1HG011543 (D.H. and M.T.), the National Institute of Mental Health of the National Institutes of Health under Award Number R01MH120295 and K12GM139185, and the Institute for the Biology of Stem Cells (IBSC) at UC Santa Cruz, the National Science Foundation under award number NSF 2034037 (M.T.), and NSF 2134955 (M.T. and D.H.), Esther A & Joseph Klingenstein Fund (to T.J.N.), Shurl and Kay Curci Foundation (T.J.N.) and the Sontag Foundation (to T.J.N.), and a gift from the William K. Bowes Jr Foundation. T.J.N. is a New York Stem Cell Foundation Robertson Neuroscience Investigator. We thank the Pacific Research Platform, the University of California Office of the President, and the University of California San Diego’s California Institute for Telecommunications and Information Technology/Qualcomm Institute. This research received software engineering support from the University of Washington’s Scientific Software Engineering Center (SSEC) supported by Schmidt Futures, as part of the Virtual Institute for Scientific Software (VISS). This project was supported by the National Institute of Neurological Disorders and Stroke, National Institutes of Health, through UCSF grant number 5R25NS070680-13 (J.P.A.). This study was also supported by NIH awards: Brain Initiative award UF1MH130700 (T.J.N.), R01NS123263 (to T.J.N.), T32HG012344 (K.V.). Its contents are solely the responsibility of the authors and do not necessarily represent the views of the NIH. The funders had no role in study design, data collection, analysis, decision to publish, or preparation of the manuscript.

## Author Contributions

M.A.T.E. and J.P.A. conceived this study, conducted the experiments, and wrote the manuscript. T.S. and D.H. supervised this work. M.A.T.E., T.V.D.M., J.G., A.S., K.V., T.G., A.S., D.F.P., A.R., and D.S. performed computational analysis of data. C.C. gave software support. A.T. constructed figures. E.F.C., T.J.N., and M.T. provided mentorship and intellectual consultation. All authors discussed the results and commented on the manuscript.

## References

1. Tatum, W. O. Mesial temporal lobe epilepsy. J. Clin. Neurophysiol. Off. Publ. Am. Electroencephalogr. Soc. 29, 356–365 (2012).

2. The Hippocampus Book. (Oxford University Press, Oxford ; New York, 2006).

3. Thom, M. Review: Hippocampal sclerosis in epilepsy: a neuropathology review. Neuropathol. Appl. Neurobiol. 40, 520–543 (2014).

4. Patrylo, P. R., Schweitzer, J. S. & Dudek, F. E. Abnormal responses to perforant path stimulation in the dentate gyrus of slices from rats with kainate-induced epilepsy and mossy fiber reorganization. Epilepsy Res. 36, 31–42 (1999).

5. Berdondini, L. et al. Active pixel sensor array for high spatio-temporal resolution electrophysiological recordings from single cell to large scale neuronal networks. Lab. Chip 9, 2644–2651 (2009).

6. Shein-Idelson, M., Pammer, L., Hemberger, M. & Laurent, G. Large-scale mapping of cortical synaptic projections with extracellular electrode arrays. Nat. Methods 14, 882–890 (2017).

7. Yuan, X. et al. Versatile live-cell activity analysis platform for characterization of neuronal dynamics at single-cell and network level. Nat. Commun. 11, 4854 (2020).

8. Andrews, J. P. et al. Multimodal evaluation of network activity and optogenetic interventions in human hippocampal slices. Nat. Neurosci. 27, 2487–2499 (2024).

9. Paulk, A. C. et al. Large-scale neural recordings with single neuron resolution using Neuropixels probes in human cortex. Nat. Neurosci. 25, 252–263 (2022).

10. Chung, J. E. et al. High-density single-unit human cortical recordings using the Neuropixels probe. Neuron 110, 2409-2421.e3 (2022).

11. Coughlin, B. et al. Modified Neuropixels probes for recording human neurophysiology in the operating room. Nat. Protoc. 18, 2927–2953 (2023).

12. Dalton, M. A., Zeidman, P., Barry, D. N., Williams, E. & Maguire, E. A. Segmenting subregions of the human hippocampus on structural magnetic resonance image scans: An illustrated tutorial. Brain Neurosci. Adv. 1, 2398212817701448 (2017).

13. Kecskés, A., Czéh, B. & Kecskés, M. Mossy cells of the dentate gyrus: Drivers or inhibitors of epileptic seizures? Biochim. Biophys. Acta Mol. Cell Res. 1869, 119279 (2022).

14. Scheibel, M. E., Crandall, P. H. & Scheibel, A. B. The hippocampal-dentate complex in temporal lobe epilepsy. A Golgi study. Epilepsia 15, 55–80 (1974).

15. Farrell, J. S.Nguyen, Q.-A. & Soltesz, I. Resolving the Micro-Macro Disconnect to Address Core Features of Seizure Networks. Neuron 101, 1016–1028 (2019).

16. Meencke, H. J., Veith, G. & Lund, S. Bilateral hippocampal sclerosis and secondary epileptogenesis. Epilepsy Res. Suppl. 12, 335–342 (1996).

17. Dudek, F. E. & Shao, L.-R. Mossy Fiber Sprouting and Recurrent Excitation: Direct Electrophysiologic Evidence and Potential Implications. Epilepsy Curr. 4, 184–187 (2004).

18. Paz, J. T. & Huguenard, J. R. Microcircuits and their interactions in epilepsy: is the focus out of focus? Nat. Neurosci. 18, 351–359 (2015).

19. Ayhan, F. et al. Resolving cellular and molecular diversity along the hippocampal anterior-to-posterior axis in humans. Neuron 109, 2091-2105.e6 (2021).

20. Govorunova, E. G. et al. Kalium channelrhodopsins are natural light-gated potassium channels that mediate optogenetic inhibition. Nat. Neurosci. 25, 967–974 (2022).

21. Buccino, A. P. et al. SpikeInterface, a unified framework for spike sorting. eLife 9, e61834 (2020).

22. Buzsáki, G., Anastassiou, C. A. & Koch, C. The origin of extracellular fields and currents--EEG, ECoG, LFP and spikes. Nat. Rev. Neurosci. 13, 407–420 (2012).

23. Radivojevic, M. et al. Tracking individual action potentials throughout mammalian axonal arbors. eLife 6, e30198 (2017).

24. Lévesque, M. & Avoli, M. The kainic acid model of temporal lobe epilepsy. Neurosci. Biobehav. Rev. 37, 2887–2899 (2013).

25. Wagenaar, D. A., Pine, J. & Potter, S. M. An extremely rich repertoire of bursting patterns during the development of cortical cultures. BMC Neurosci. 7, 11 (2006).

26. King, D. & Spencer, S. Invasive electroencephalography in mesial temporal lobe epilepsy. J. Clin. Neurophysiol. Off. Publ. Am. Electroencephalogr. Soc. 12, 32–45 (1995).

27. Jones, R. S. G., da Silva, A. B., Whittaker, R. G., Woodhall, G. L. & Cunningham, M. O. Human brain slices for epilepsy research: Pitfalls, solutions and future challenges. J. Neurosci. Methods 260, 221–232 (2016).

28. Kitchigina, V. F. Alterations of Coherent Theta and Gamma Network Oscillations as an Early Biomarker of Temporal Lobe Epilepsy and Alzheimer’s Disease. Front. Integr. Neurosci. 12, 36 (2018).

29. Lubenov, E. V. & Siapas, A. G. Hippocampal theta oscillations are travelling waves. Nature 459, 534–539 (2009).

30. Zhang, H. & Jacobs, J. Traveling Theta Waves in the Human Hippocampus. J. Neurosci. Off. J. Soc. Neurosci. 35, 12477–12487 (2015).

31. Diamond, J. M. et al. Interictal discharges in the human brain are travelling waves arising from an epileptogenic source. Brain J. Neurol. 146, 1903–1915 (2023).

32. Katzner, S. et al. Local Origin of Field Potentials in Visual Cortex. Neuron 61, 35–41 (2009).

33. Whittingstall, K. & Logothetis, N. K. Frequency-Band Coupling in Surface EEG Reflects Spiking Activity in Monkey Visual Cortex. Neuron 64, 281–289 (2009).

34. Siapas, A. G., Lubenov, E. V. & Wilson, M. A. Prefrontal phase locking to hippocampal theta oscillations. Neuron 46, 141–151 (2005).

35. Rutishauser, U., Ross, I. B., Mamelak, A. N. & Schuman, E. M. Human memory strength is predicted by theta-frequency phase-locking of single neurons. Nature 464, 903–907 (2010).

36. Schonhaut, D. R. et al. MTL neurons phase-lock to human hippocampal theta. eLife 13, e85753 (2024).

37. Anastassiou, C. A., Perin, R., Markram, H. & Koch, C. Ephaptic coupling of cortical neurons. Nat. Neurosci. 14, 217–223 (2011).

38. Cutts, C. S. & Eglen, S. J. Detecting Pairwise Correlations in Spike Trains: An Objective Comparison of Methods and Application to the Study of Retinal Waves. J. Neurosci. 34, 14288–14303 (2014).

39. Buchin, A. et al. Multi-modal characterization and simulation of human epileptic circuitry. Cell Rep. 41, 111873 (2022).

40. Sharf, T. et al. Functional neuronal circuitry and oscillatory dynamics in human brain organoids. Nat. Commun. 13, 4403 (2022).

41. Koch, C., Rapp, M. & Segev, I. A brief history of time (constants). Cereb. Cortex N. Y. N 1991 6, 93–101 (1996).

42. English, D. F. et al. Pyramidal Cell-Interneuron Circuit Architecture and Dynamics in Hippocampal Networks. Neuron 96, 505-520.e7 (2017).

43. Barthó, P. et al. Characterization of Neocortical Principal Cells and Interneurons by Network Interactions and Extracellular Features. J. Neurophysiol. 92, 600–608 (2004).

44. Shin, D. et al. High-Complexity Barcoded Rabies Virus for Scalable Circuit Mapping Using Single-Cell and Single-Nucleus Sequencing. 2024.10.01.616167 Preprint at 10.1101/2024.10.01.616167 (2024).

45. King-Stephens, D. What Is the Best Target for Ablation of Mesial Temporal Lobe Epilepsy? Epilepsy Curr. 19, 313 (2019).

46. Aguila, C. A. et al. Mesial-to-lateral patterns of epileptiform activity identify the seizure onset zone in mesial temporal lobe epilepsy. 2024.10.29.24316309 Preprint at 10.1101/2024.10.29.24316309 (2024).

47. Ting, J. T. et al. A robust ex vivo experimental platform for molecular-genetic dissection of adult human neocortical cell types and circuits. Sci. Rep. 8, 8407 (2018).

48. Krook-Magnuson, E., Armstrong, C., Oijala, M. & Soltesz, I. On-demand optogenetic control of spontaneous seizures in temporal lobe epilepsy. Nat. Commun. 4, 1376 (2013).

49. Huberfeld, G. et al. Glutamatergic pre-ictal discharges emerge at the transition to seizure in human epilepsy. Nat. Neurosci. 14, 627–634 (2011).

50. Müller, J. et al. High-resolution CMOS MEA platform to study neurons at subcellular, cellular, and network levels. Lab. Chip 15, 2767–2780 (2015).

51. Pachitariu, M., Steinmetz, N., Kadir, S., Carandini, M. & D, H. K. Kilosort: realtime spike-sorting for extracellular electrophysiology with hundreds of channels. 061481 Preprint at 10.1101/061481 (2016).

52. Smarr, L. et al. The Pacific Research Platform: Making High-Speed Networking a Reality for the Scientist. in Proceedings of the Practice and Experience on Advanced Research Computing 1–8 (Association for Computing Machinery, New York, NY, USA, 2018). doi:10.1145/3219104.3219108.

53. Rossant, C. et al. Spike sorting for large, dense electrode arrays. Nat. Neurosci. 19, 634–641 (2016).

54. Guth, T. A. et al. Theta-phase locking of single neurons during human spatial memory. bioRxiv 2024.06.20.599841 (2024) doi:10.1101/2024.06.20.599841.

55. Santhakumar, V., Aradi, I. & Soltesz, I. Role of mossy fiber sprouting and mossy cell loss in hyperexcitability: a network model of the dentate gyrus incorporating cell types and axonal topography. J. Neurophysiol. 93, 437–453 (2005).

