## Supplementary Information for "Pathological microcircuits and epileptiform events in patient hippocampal slices"

**This PDF Includes:**

Extended Data Fig. 1 to 12

Extended Data Table 1 to 4

Video 1 to 4

| Recording | Slice | Patient ID | Region | Kainic Acid | Recurrent Circuit | Seizure-Like Behaviour | Recording Length (minutes) | Neuron Count |
|---|---|---|---|---|---|---|---|---|
| 1 | S1 | A | Dentate Gyrus | Yes | Yes | Yes | 6 | 212 |
| 2 | | | | No | Yes | No | 5 | 249 |
| 3 | S2 | B | Dentate Gyrus | Yes | Yes | Yes | 8.5 | 180 |
| 4 | | | | No | Yes | No | 5 | 51 |
| 5 | S3 | C | Dentate Gyrus | No | No | No | 7 | 87 |
| 6 | S4 | D | Dentate Gyrus | Yes | No | No | 2.2 | 109 |
| 7 | S5 | E | CA1 | No | No | No | 6.4 | 136 |
| 8 | S6 | F | CA1 | No | No | No | 1.8 | 100 |

**Extended Data Table 1 |Summary of electrophysiology recordings.** Summary of the eight electrophysiology recordings used for analysis. Data comes from 6 hippocampal slices, each from a different patient. Four slices are from the dentate gyrus and two are from CA1. Two of the dentate gyrus slices exhibited seizure-like behavior. These slices (S1 and S2) had two recordings each, a baseline recording and a recording after the administration of kainic acid.

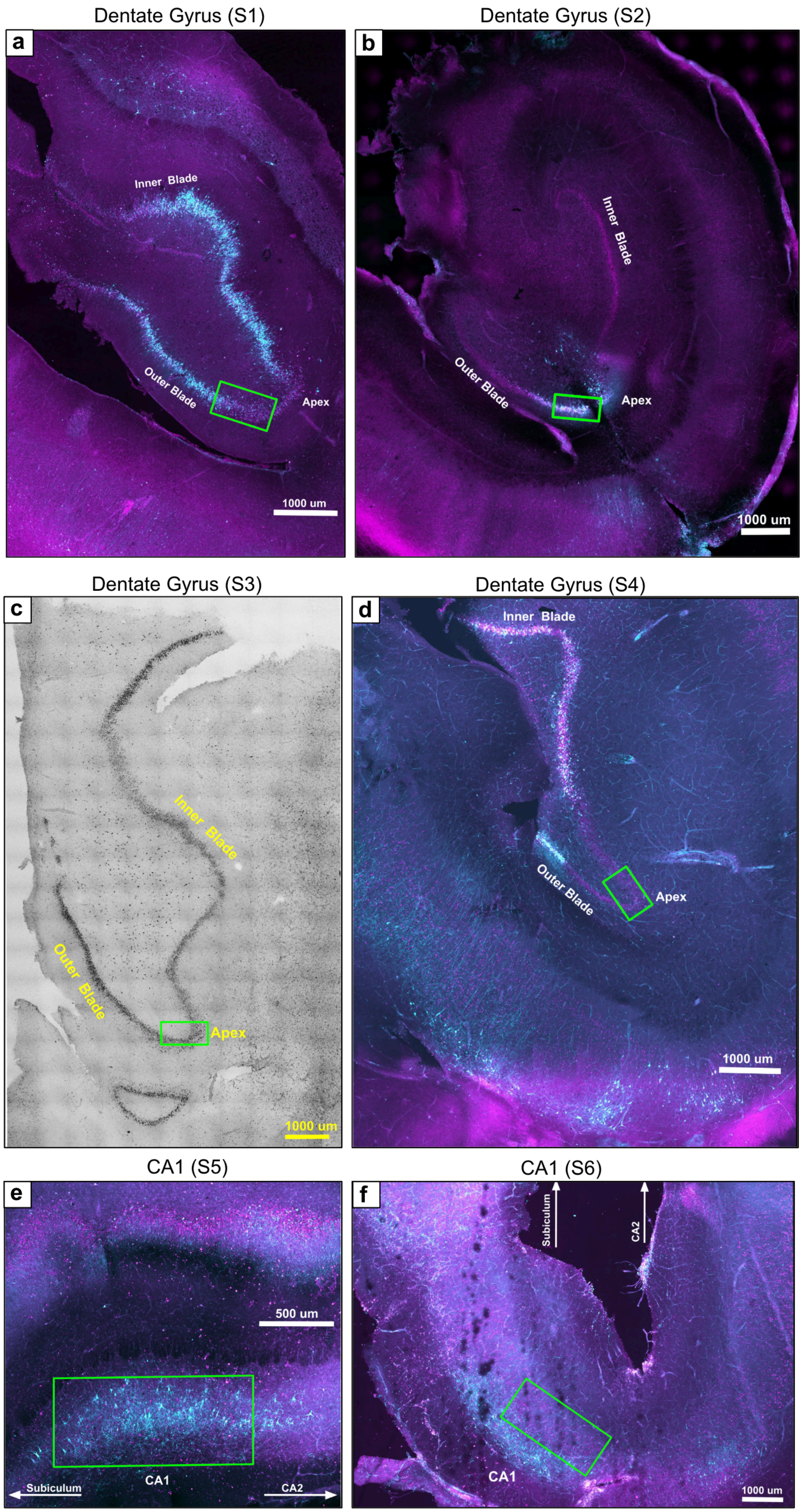


**Extended Data Fig. 1 | Histology stains for all slices.** NeuN (magenta) and eYFP (cyan) immunohistochemistry of slices transduced with AAV9-CAMK2A-HcKCR1-eYFP, with cyan representing CAMK2A expressing cells. For each slice, a green box highlights the region where neural activity was recorded. Slices S1-S4 are from the dentate gyrus. **a-b,** S1-S2 had recordings taken from the outer blade, proximal to the apex. **c-d,** S3-S4 had recordings taken from the inner apex. **e-f,** Slices S5-S6 contain recordings from CA1.

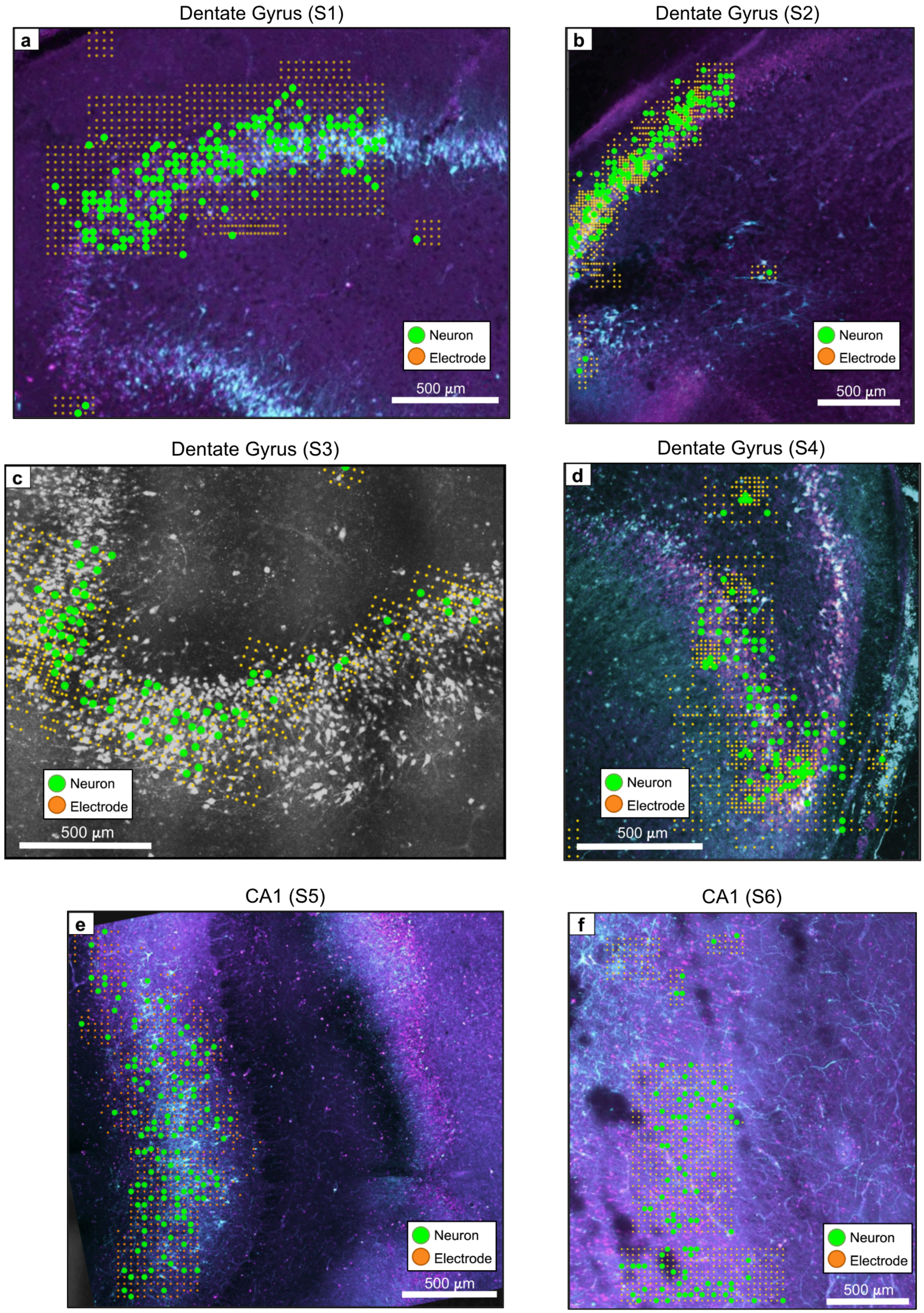


**Extended Data Fig. 2 | Spatial map of electrodes and neurons by slice.** Spatial map of recording electrodes and spike sorted neural units for all six slices where recordings were taken. Orange dots are electrodes, and green circles are neurons. **a-b,** The recordings from slices S1 and S2 were taken from approximately the same region of the dentate gyrus. **c-d,** Slices S3 and S4 contain recordings from the apex of the dentate gyrus. **e-f,** Slices S5 and S6 were recorded within the CA1 region.

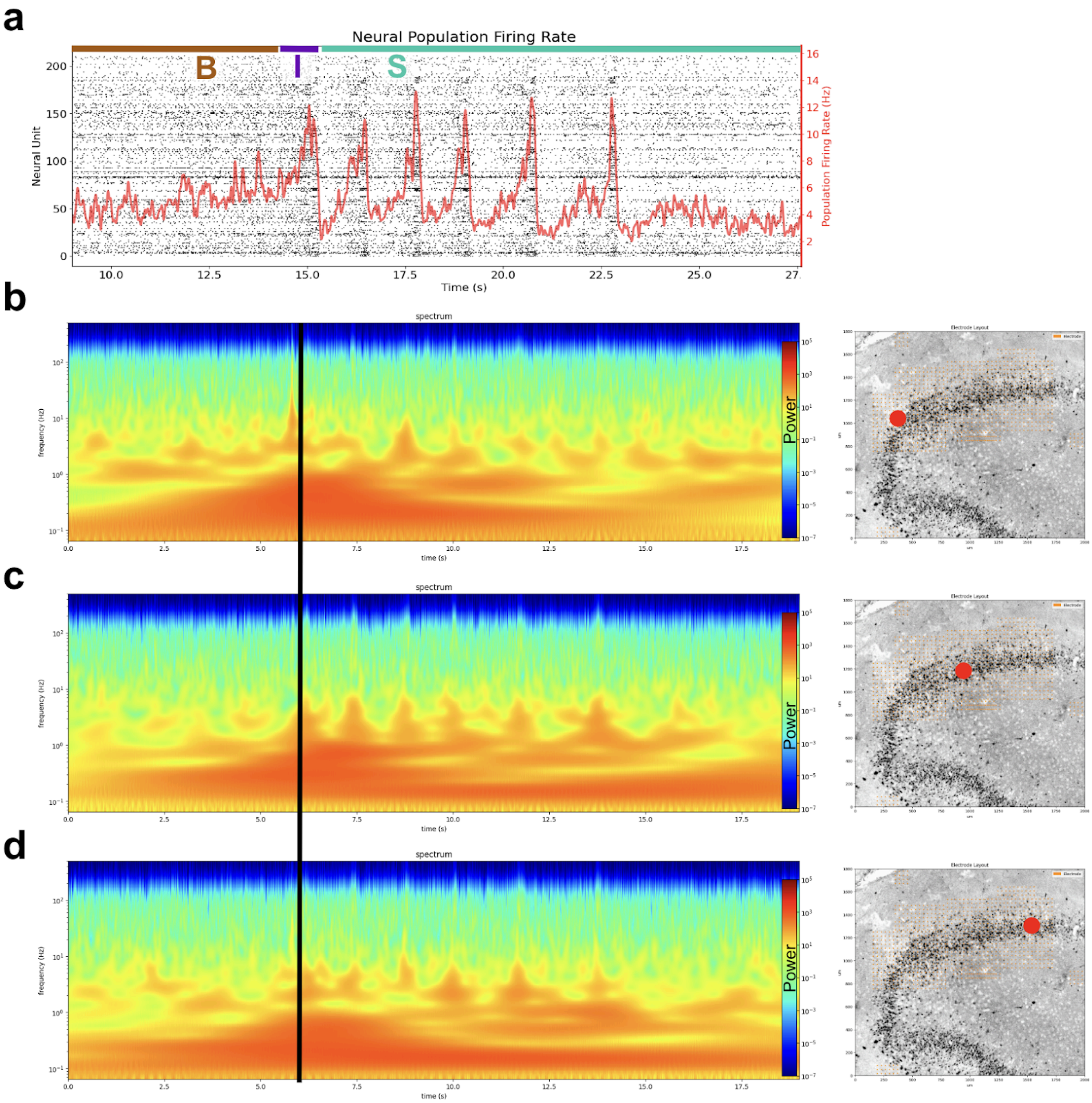


**Extended Data Fig. 3 | Spectrogram results are invariant to electrode location. a,** A plot of neural activity from the first superburst of the recording. The baseline, initiation, and seizure phases are labeled B, I, and S respectively. **b-c,** (left) Spectrograms from three different electrodes are displayed. (right) For each spectrogram, the red circle on the histology image displays the location of the electrode used for the spectrogram. All three spectrograms display a large upwelling in power in the delta frequency near the initiation phase of the recording (vertical black line) as well as theta activity aligned with bursts.

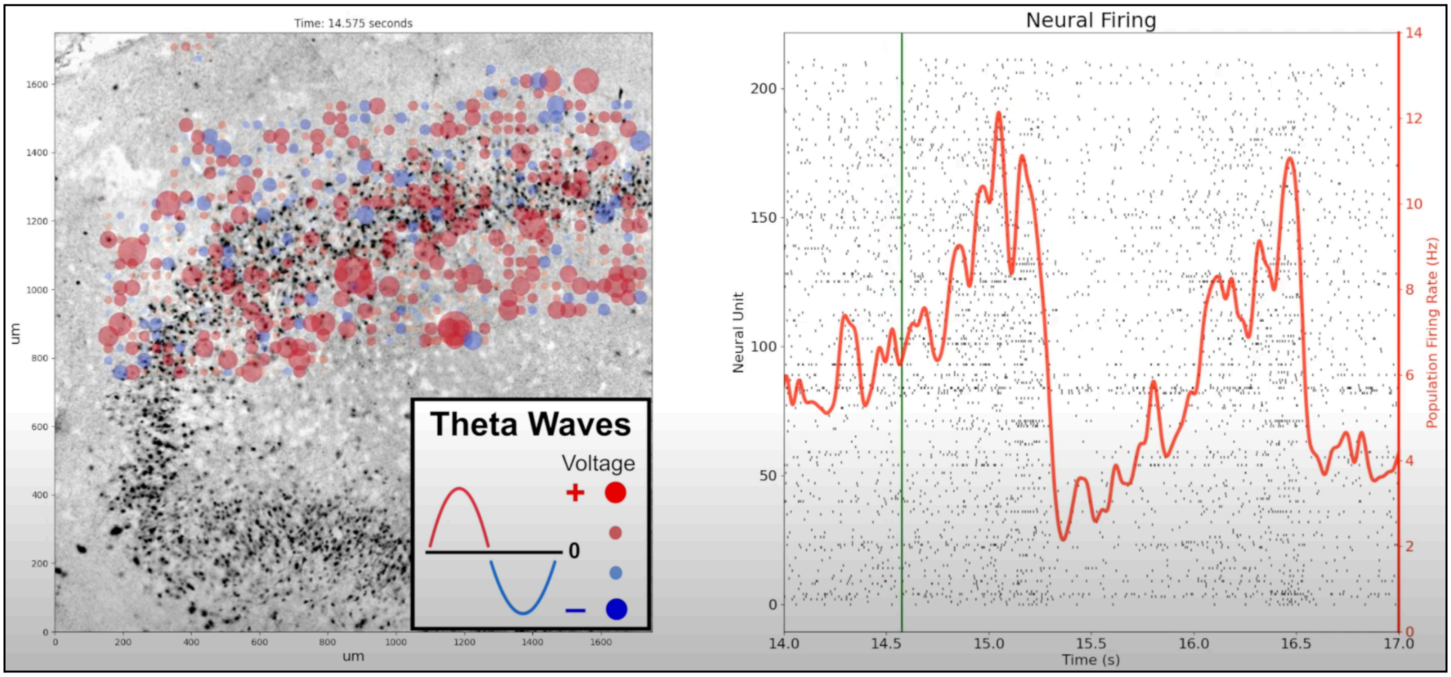


https://youtu.be/sm3gKk-1Og4

**Video 1 | Theta wave propagations at onset of epileptiform activity.** Video of theta wave behavior during the initiation of epileptiform activity. The video considers two slices, both with recordings taken from the dentate gyrus (S1, S2). For both slices, theta behavior follows three phases: baseline line, initiation, and seizure stage.

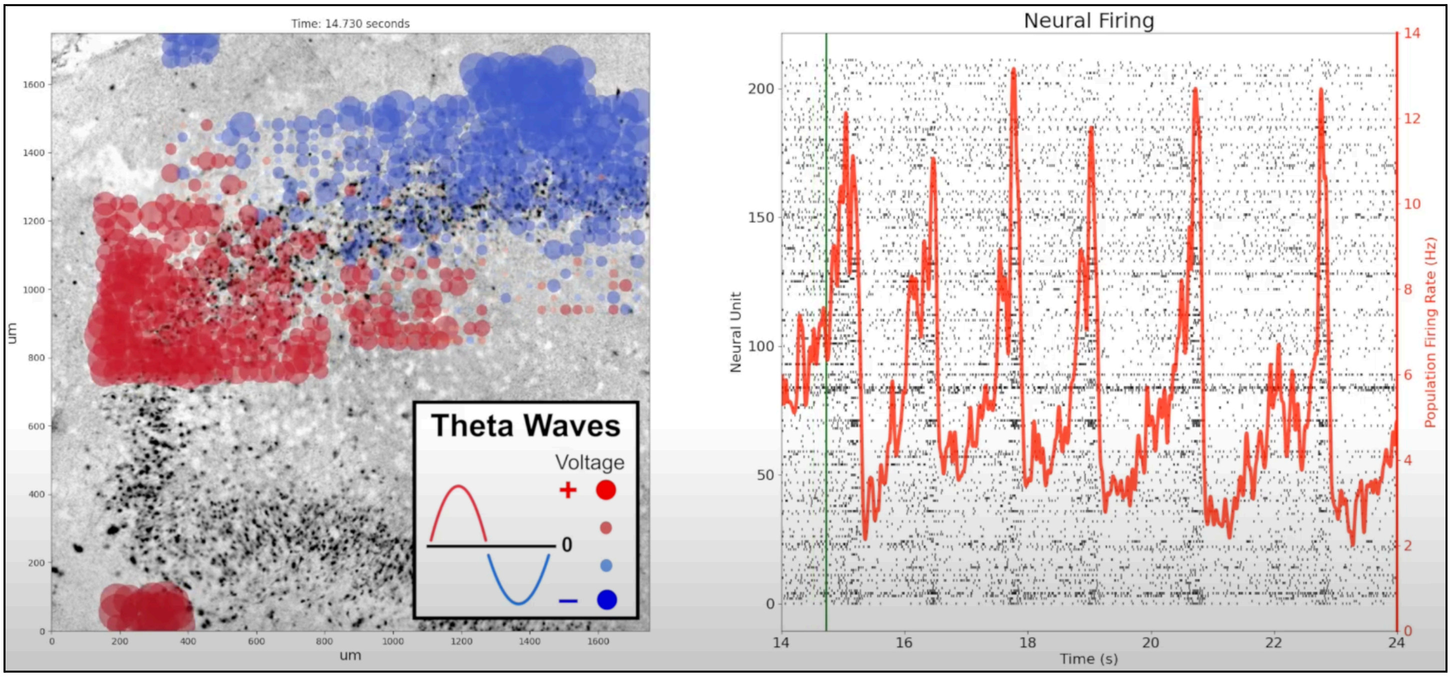


**https://www.youtube.com/watch?v=mqjJhCuNR_A**

**Video 2 | Theta wave propagations from the first superburst.** Video of theta wave behavior across the entirety of the first superburst during epileptiform activity. The video is from the slice used in Fig. 1 and Video 1 (S1). Notice that the theta behavior observed at the initiation of epileptiform activity is not replicated anywhere else in the recording.

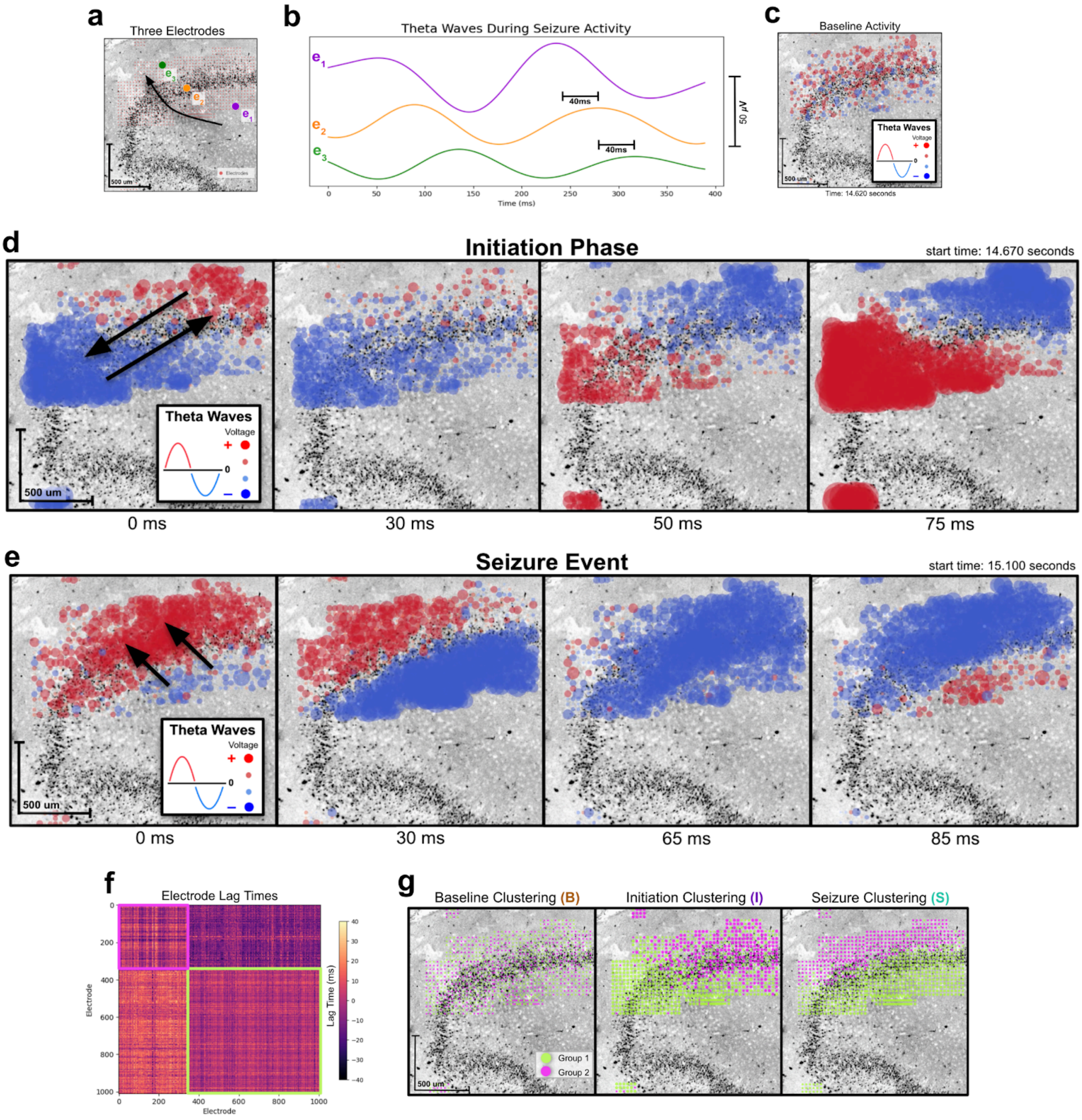


**Extended Data Fig. 4 | Seizure initiation phase drives coherent theta wave propagations. a,** The spatial location of three electrodes is displayed (e1,e2,e3). A black arrow represents the direction that theta waves propagate during seizure-like activity. **b,** Theta waves from (e1,e2,e3) taken from a 400ms time interval of seizure-like activity. There is a 40ms delay between theta waves as they propagate from  e1 to e2, and from  e2 to e3. **c,** A spatial plot of theta wave activity during baseline activity. No discernible pattern was observed. **d,** A timelapse of theta wave plots from the initiation phase over the course of 75ms. Black arrows in the left-most plot display the direction of propagations. Propagations form a standing wave that oscillates across the length of the granule cell layer. **e,** A timelapse from the seizure phase over the course of 85ms. Propagations form a rolling wave, moving from the hilar aspect to the outer aspect. **f,** A matrix of the lag times (ms) between theta waves for all electrode pairs. Pink and green squares present electrode clusters produced using the k-means algorithm. **g,** Spatial plots of electrode groups during baseline, initiation, and seizure phase. Groupings are based on the k-means clustering described in plot f. Clusterings resemble theta activity seen in the corresponding phase.

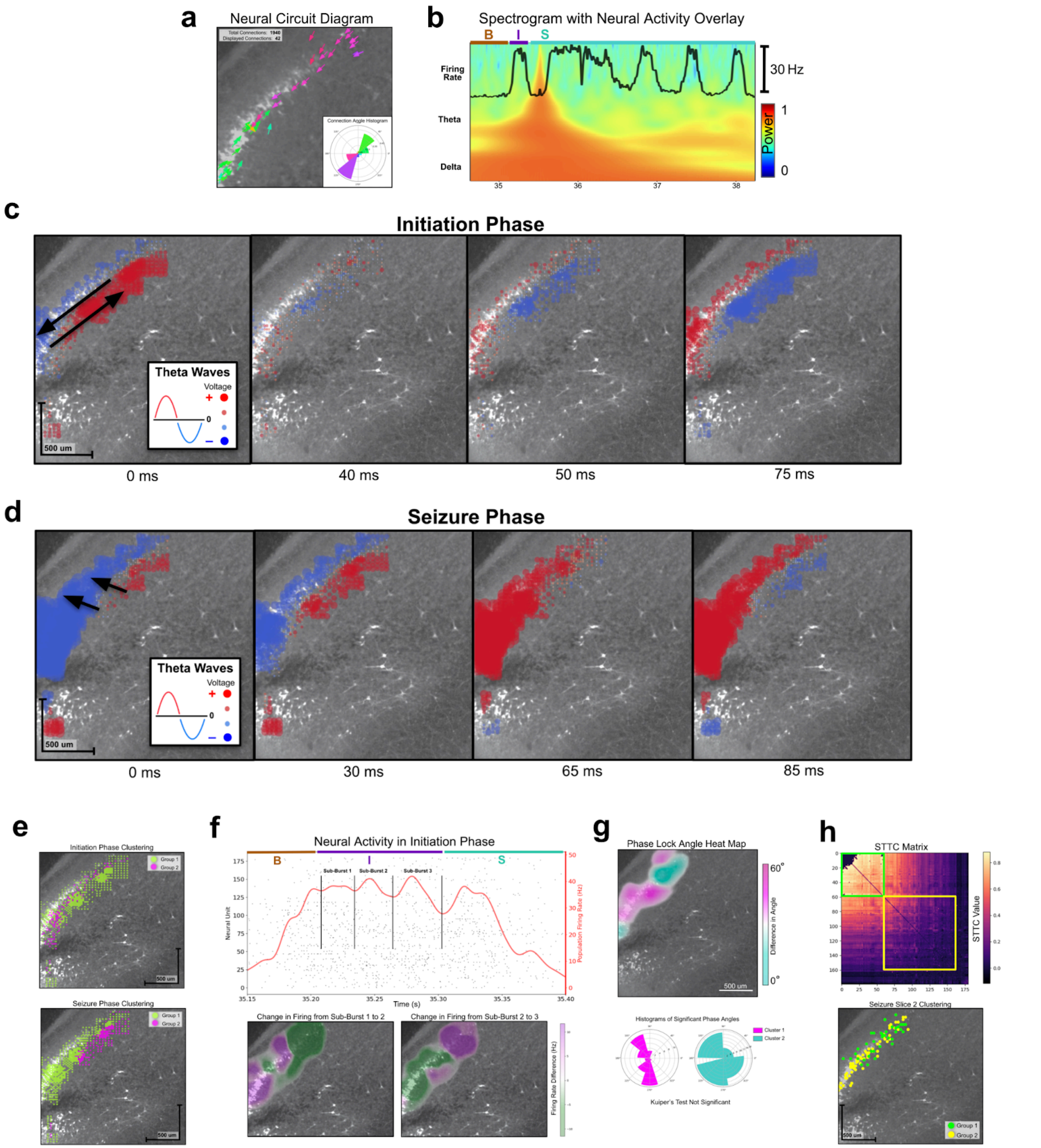


**Extended Data Fig. 5 | Epileptiform activity from second slice (S2) is consistent with first (S1). a,** The neural circuit diagram from S2. The connection angle histogram displays a bimodal circuit (bottom-right). **b,** A spectrogram of delta and theta activity with the neural firing rate overlaid on top of it. The recording is divided into baseline (B), Initiation (I), and Seizure (S) phases. A large upwelling in delta and theta activity occurs near the initiation phase. **c,** A timelapse of theta wave plots from the initiation phase over the course of 75ms. Black arrows in the left-most plot display the direction of propagations. Propagations form a standing wave. **d,** A timelapse from the seizure phase over the course of 85ms. Propagations form a rolling wave, moving from the hilar aspect to the outer aspect. **e,** Spatial plot of electrode clusterings during the initiation and seizure phases. **f,** (Top) Neural activity during the initiation phase (I) is divided into three sub-bursts. (bottom) Spatial plots of the change in neural firing from sub-burst to sub-burst show bimodal recurrent behavior. **g,** (Top) A heatmap of the average difference in angles between all significantly phase locked units. (Bottom) Histograms of phase angles from the clusters seen in the heatmap. **h,** (Top) STTC matrix with units organized based on an agglomerative hierarchical clustering algorithm. Green and yellow squares indicate groupings observed from clustering. (Bottom) Spatial plot of neurons based on their grouping from the STTC matrix. Green and yellow groups correspond to the squares on the STTC matrix.

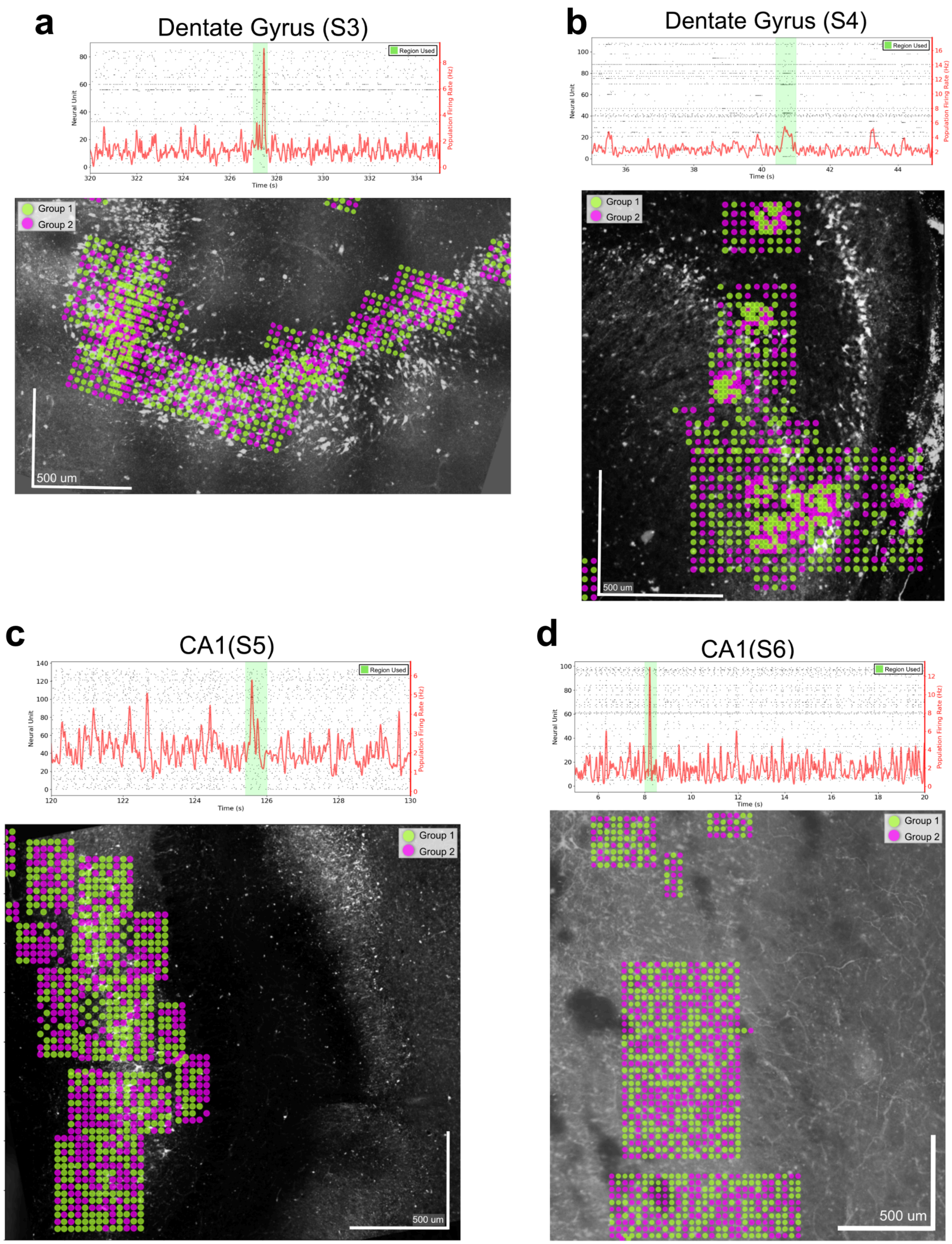


**Extended Data Fig. 6 | Slices without seizure behavior have no spatial clustering in their electrodes.** Spatial plots of electrode groupings for slices with no epileptiform activity (S3-S6). Unlike the epileptiform slices, no pattern was observed in these clusterings. For all slices, the top plot depicts the region of the recording used for clustering (green). Recordings were sampled at points that contained higher than average burst-like activity. **a-b,** Plots from the dentate gyrus (S3-S4). **c-d,** Plots from the CA1 (S5-S6).

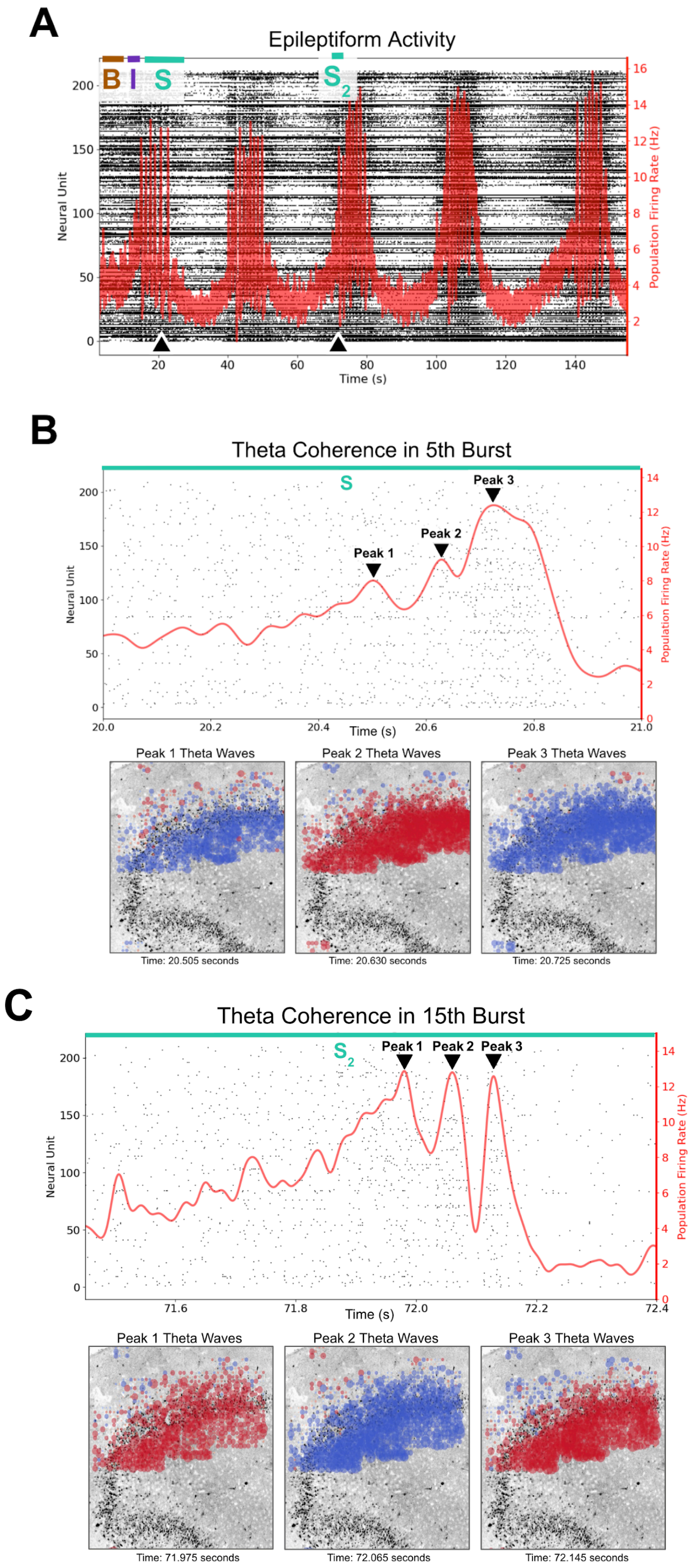


**Extended Data Fig. 7 | Theta coherence occurs across the seizure phase of the recording. a,** Neural activity plot during epileptiform activity for slice S1. Black triangles at the base of the recording indicate the locations of the two bursts shown in plots b-c. **b-c,** Coherence between propagating theta waves and the population level firing activity within the first and third superburst. (Top) In the burst's neural activity plot, black triangles indicate the locations of sub-burst peaks. (Bottom) Theta activity plots corresponding to the peaks. The timing of wave propagations align with the peaks. Note, Video 3 displays wave propagations for bursts in b and c.

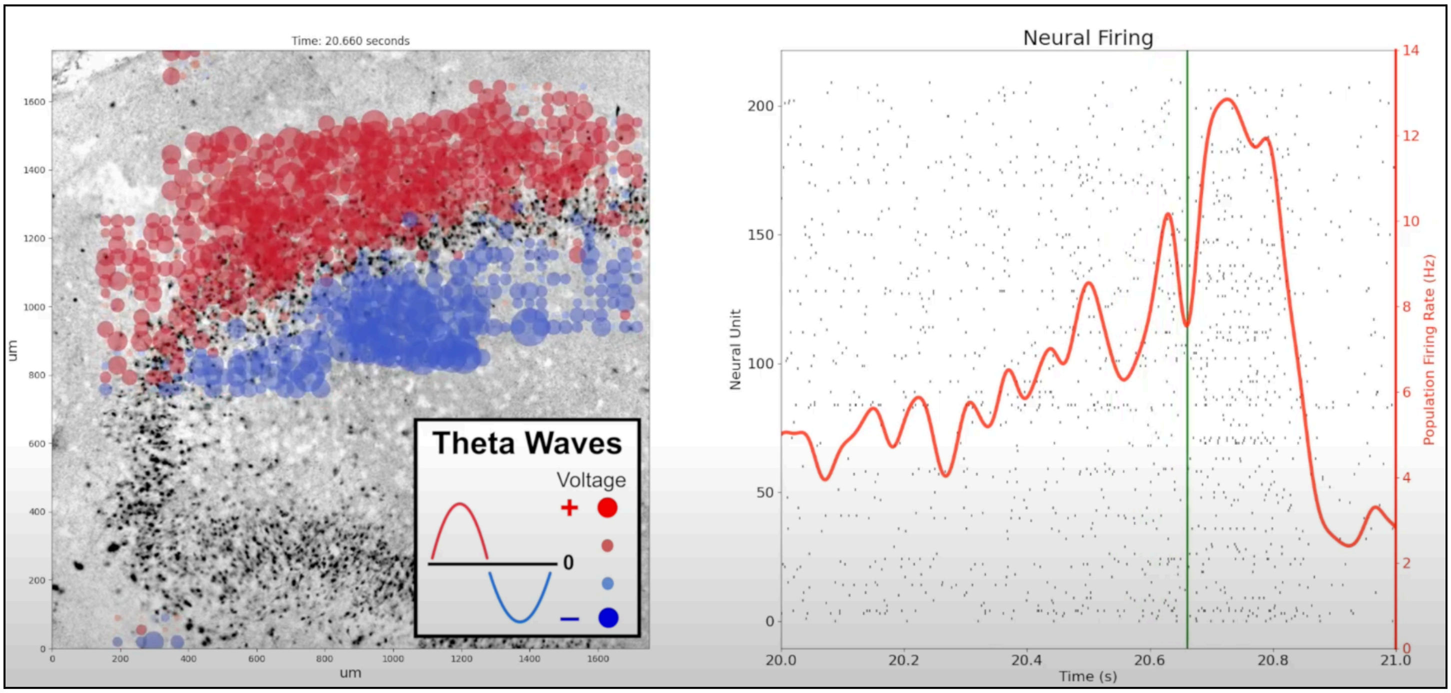


https://www.youtube.com/watch?v=I_5XFMt1Lh0

**Video 3 | Theta wave coherence in seizure-like bursts.** Video of theta wave behavior during epileptiform activity from the 5th and 15th burst of slice S1. These bursts are shown in Extended Data Fig. 7. In both bursts, the timing of the theta wave propagations align with the the burst peaks.

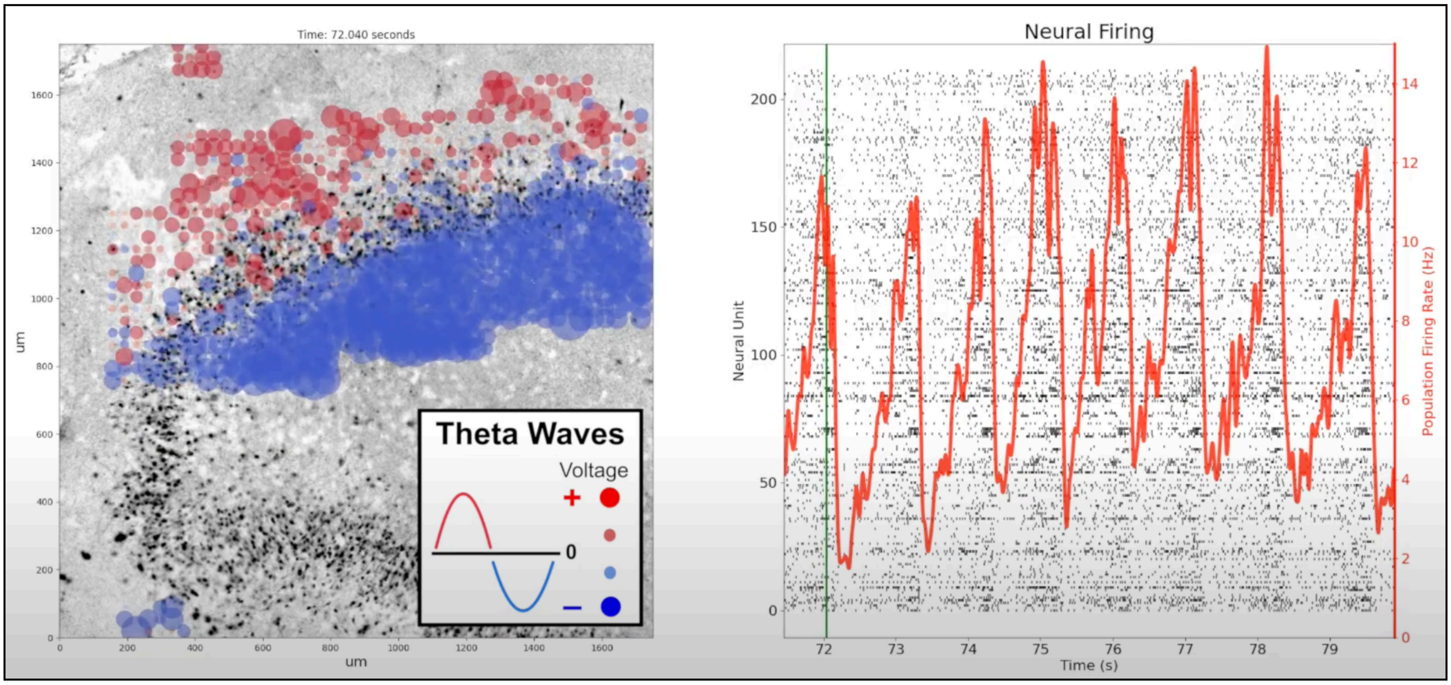


https://www.youtube.com/watch?v=1AIXzt8AVOU

**Video 4 | Theta Wave Propagations in third superburst.** Video of theta wave behavior across the entirety of the third superburst during epileptiform activity in slice S1. Unlike the first superburst, theta activity never displays an initiation phase.

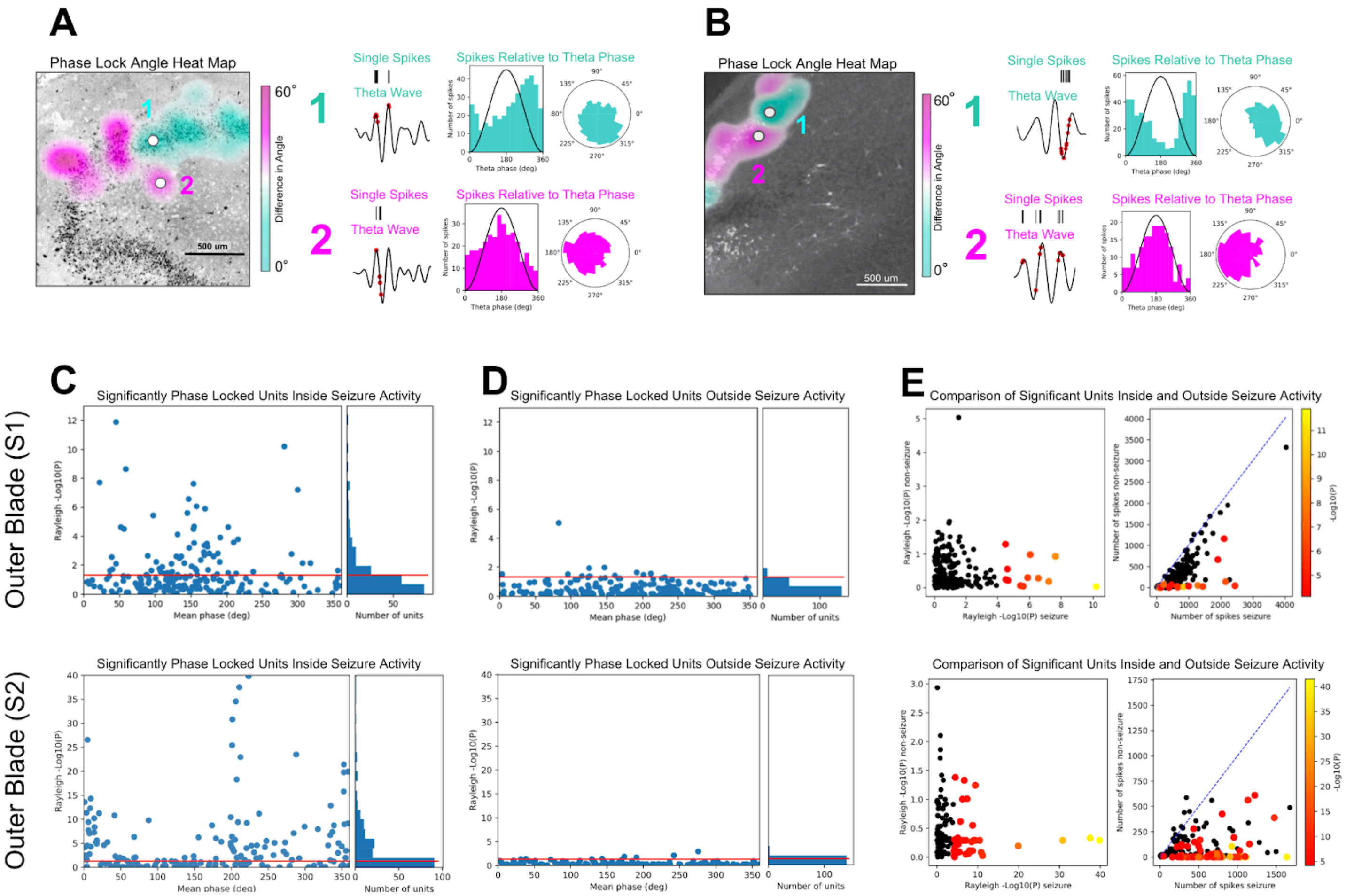


**Extended Data Fig. 8 | Theta phase locking is more prevalent during seizure-like behavior. a-b,** The spatial locations of the two neural units, one from each cluster of phase locked units for epileptiform slices S1 and S2. We show phase angle histogram for each unit. **c-d** (left) Plots display the significance of phase locked units by their phase angle. Neurons above the red horizontal line are statistically significant (Rayleigh p-value<0.05) (right) Histogram displays to total aggregate number of units by significance. Plot C is during seizure activity, and Plot D is outside seizure activity. In S1 and S2, there are considerably more phase locked units during seizure-like behavior. **e,** (left) A scatter comparing the significance of phase locked units from within and outside seizure behavior. Seizure behavior exhibits more phase locking. (right) The figure on the right shows that the majority of the units that are phase locked during seizure events are not active during non-seizure events. Such units are only active during seizure-like behavior.

| Recording | Slice | Region | Kainic Acid | Recurrent Circuit | Seizure-Like Behaviour | Total Neurons | Significant Neurons | Percent Significant | |
|---|---|---|---|---|---|---|---|---|---|
| 1 | S1 | Dentate Gryus | Yes | Yes | Yes | 212 | 70 | 33% | * 5% Outside Seizure |
| 2 | | | No | Yes | No | 249 | 14 | 6% | |
| 3 | S2 | Dentate Gryus | Yes | Yes | Yes | 180 | 101 | 56% | * 4% Oustide Seizure |
| 4 | | | No | Yes | No | 51 | 2 | 4% | |
| 5 | S3 | Dentate Gyrus | No | No | No | 87 | 8 | 9% | |
| 6 | S4 | Dentate Gyrus | Yes | No | No | 109 | 6 | 6% | |
| 7 | S5 | CA1 | No | No | No | 136 | 8 | 6% | |
| 8 | S6 | CA1 | No | No | No | 100 | 1 | 1% | |

**Extended Data Table 2 | Theta phase locking is more prevalent during epileptiform activity.** Table displaying the number of units significantly phase locked to the theta frequency for each recording (Rayleigh p-value<0.05). The recordings with epileptiform activity are denoted in the column “Seizure-Like Behavior”. The last column displays the total percentage of phase locked units. During seizure-like activity, on average, 45% of neurons per recording are phase locked. This is considerably higher than the 5% average seen in recordings with no seizure-like activity. In the recordings that expressed seizure-like behavior, average phase locking was considerably higher in the portions of the recording with epileptiform activity, compared to the portions without (see *).

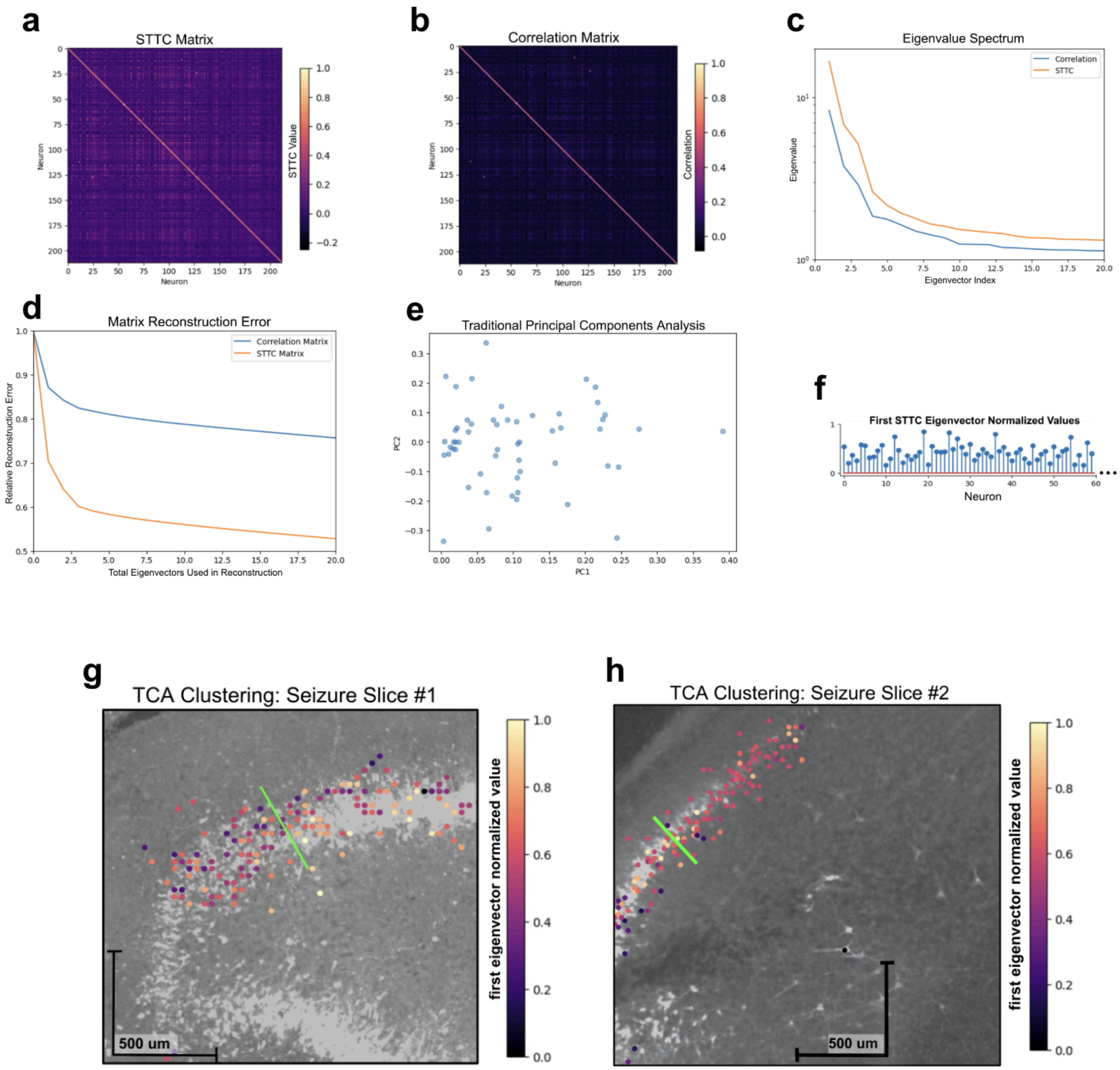


**Extended Data Fig. 9 | STTC eigendecomposition recapitulates spatial clustering in neurons. a-b,** The spike time tiling coefficient (STTC) matrix and the correlation matrix from slice S1. The STTC matrix is less sparsely populated than the correlation matrix. **c,** The first 20 eigenvalues, ranked by value, from the STTC (orange) and correlation (blue) matrices. Eigenvalues are larger for the STTC matrix. **d,** The matrix reconstruction error from the STTC (orange) and correlation (blue) matrices, measuring the extent to which eigenvectors reconstruct the matrix. The reconstruction error is lower for the STTC. **e,** Traditional PCA scatter plot comparing the first two eigenvectors. **f,** The first eigenvector from the STTC matrix, with the eigenvector value of each neuron (x-axis) displayed (y-axis). **g-h,** The first eigenvector spatially mapped to neurons for both epileptiform recordings (S1 and S2). Both recordings express a gradient moving across the granule cell layer.

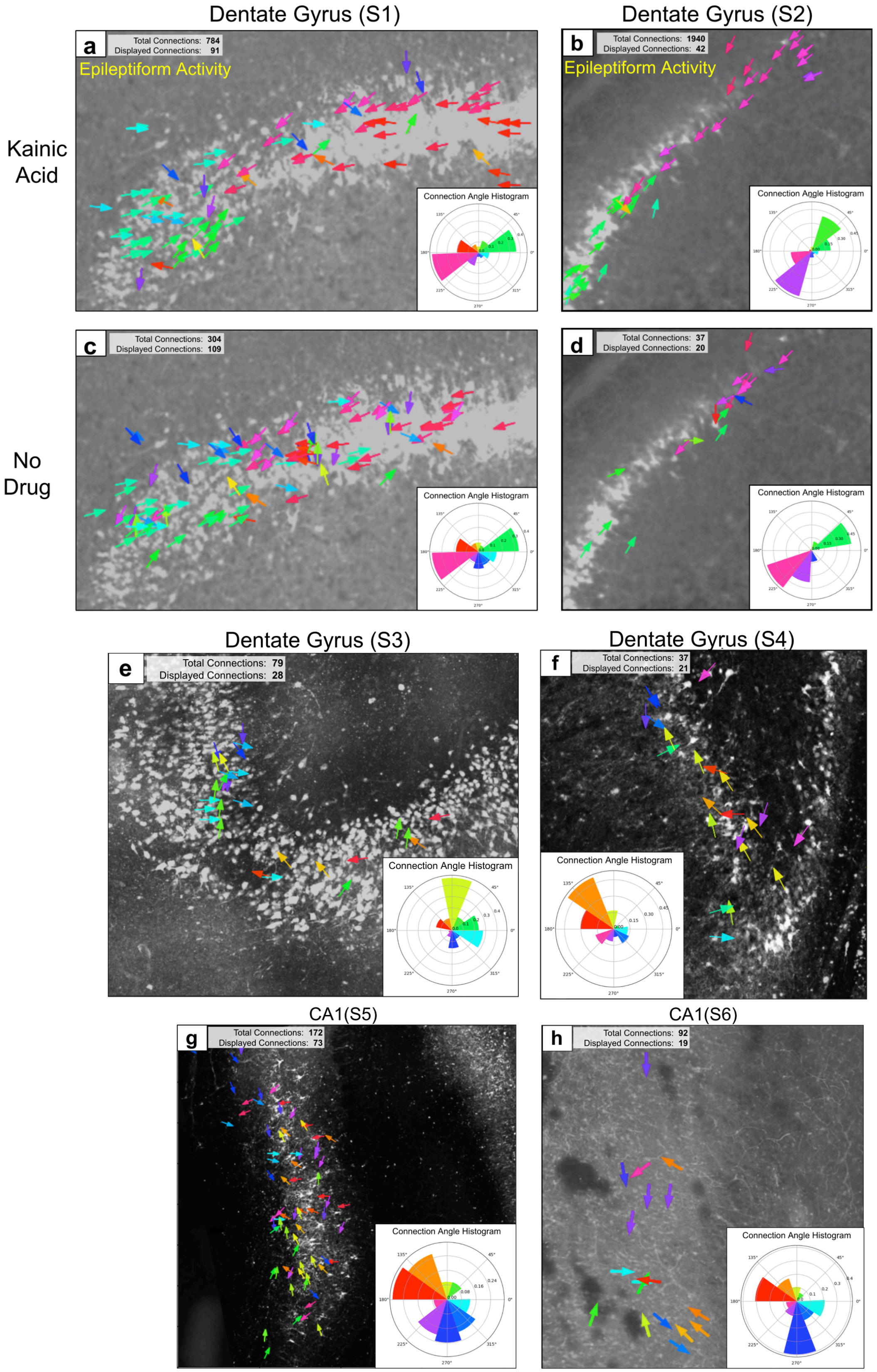


**Extended Data Fig. 10 | Neural circuit diagrams by recording.** Neural circuit diagrams for every recording. In every plot, vector arrows are colored based on the angle that spikes propagate. In the top-left, we indicate the total number of pairwise correlations and the number of correlations that are displayed. At the bottom, the pairwise correlation angle histogram depicts the angular frequencies from all spike propagation events. **a-d,** Diagrams from slices S1 and S2, both taken from the dentate gyrus. Diagrams a-b are from epileptiform recordings, after the administration of kainic acid. Plots c-d are from baseline recordings before kainic acid was administered. All plots display a bipolar recurrent circuit across the granular cell layer. **e-f,** Diagrams from the dentate gyrus (S3 and S4) display a neural circuit with spike signals propagating from the apex toward the inner blade. **g-h,** Diagrams from CA1 (S5 and S6) have neural circuits with spike signals that propagate down the perforant pathway toward the subiculum, as well as another signal that propagates from the dentate gyral aspect toward the entorhinal aspect.

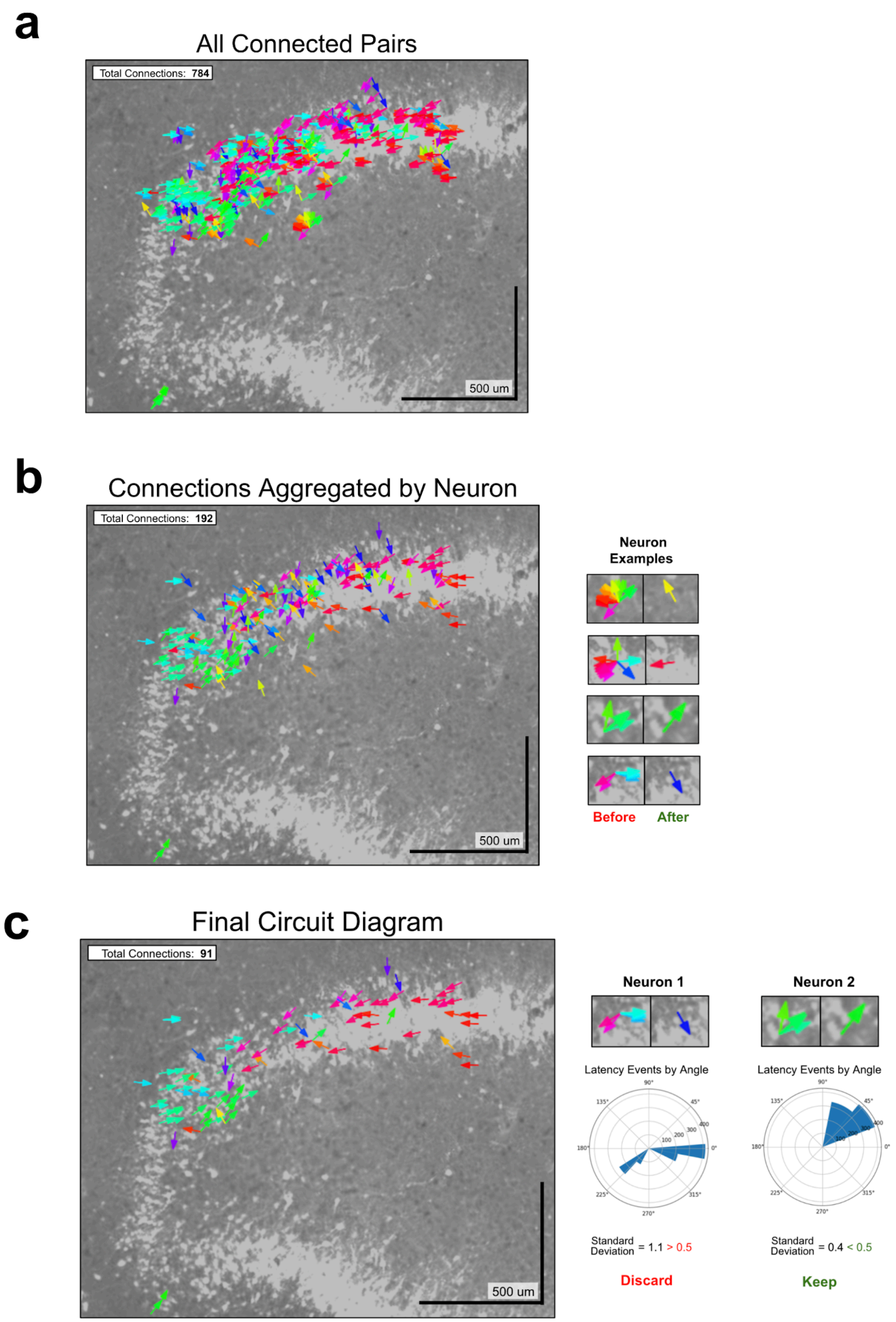


**Extended Data Fig. 11 | Schematic for curating displayed circuit connections. a,** A circuit diagram of all 784 significant connections found from slice S1. The number of arrows makes the plot less interpretable. **b,** (left) Circuit diagram after connections are aggregated based on the neuron propagating the signal. (right) Examples of connections before and after they are aggregated. The bottom example depicts why aggregation leads to a disproportionately large number of blue arrows. **c,** (left) The final circuit diagram for S1. (right) Two examples illustrate how the standard deviation of aggregated connections is used to remove non-representative arrows.

| Recording | Slice | Region | Kainic Acid | Recurrent Circuit | T-test P-value |
|---|---|---|---|---|---|
| 1 | S1 | Dentate Gyrus | Yes | Yes | 9.7e-20 |
| 2 | | | No | Yes | 8.6e-13 |
| 3 | S2 | Dentate Gyrus | Yes | Yes | 1.6e-68 |
| 4 | | | No | Yes | 1.0e-6 |
| 5 | S3 | Dentate Gyrus | No | No | 0.12 |
| 6 | S4 | Dentate Gyrus | Yes | No | 0.37 |
| 7 | S5 | CA1 | No | No | 0.00042 |
| 8 | S6 | CA1 | No | No | 0.0011 |

**Extended Data Table 3 | Results from the circuit geometry test by recording.** Results from the circuit geometry test for all recordings. The last column displays the resulting p-value of the test by recording. If the results of the circuit connection diagrams were due to bias caused by the geometry of the neurons, we would expect all slices to have approximately the same p-value. The recordings with pathological recurrent circuits have p-values orders of magnitudes lower than the other recordings.

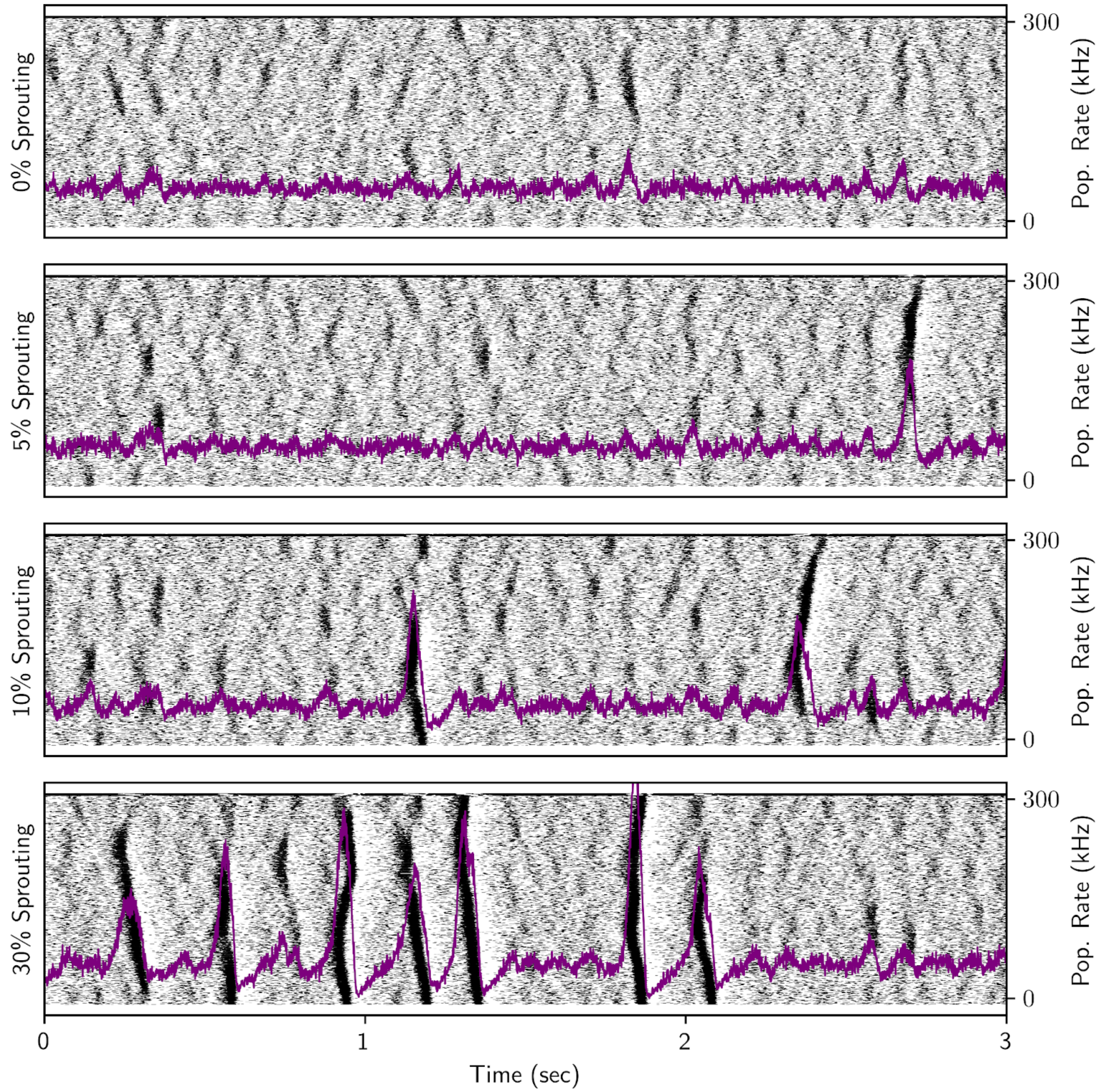


**Extended Data Fig. 12 | Simulation of pathological interconnectivity in the dentate gyrus.** Simulations of dentate gyrus hyperexcitability caused by mossy fiber sprouting. The simulations were conducted four times with varying levels of pathological recurrent connectivity in the dentate gyrus. The spike raster (black) is overlaid with the population firing rate (purple). The simulation was performed with 1000 granule cells. As interconnectivity between cells increased, more seizure-like behavior was observed.

.

| Slice | Age | Sex | ILAE Sclerosis Score | Neurons | Perturbations |
|---|---|---|---|---|---|
| S1 | 52 | F | type 1 | 212 | 0-mg +/- KA |
| S2 | 35 | M | No Hs | 180 | 0-mg +/- KA |
| S3 | 20 | M | type 3 | 87 | none |
| S4 | 46 | F | type 2 | 109 | 0-mg +/- KA |
| S5 | 40 | F | type 3 | 136 | none |
| S6 | 46 | F | No Hs | 100 | none |

**Extended Data Table 4 | Patient information.** Characteristics of patients from which hippocampal slices were excised. Age in years and sex (M=male, F=female) are shown. The International League Against Epilepsy (ILAE) classification system was used for scoring the severity of hippocampal sclerosis within the slides based on their histology, with severity going from no sclerosis (No Hs) to type 3. Perturbations refer to changes to media during the course of experiments, with 0-mg +/- KA denoting the use of zero magnesium media with and the addition of kainic acid.
